# Eight years of social and individual learning jointly drive ecological competence in orangutans

**DOI:** 10.1101/2025.08.27.672553

**Authors:** T Revathe, Marie-Theres Weidling, Sri Suci Utami-Atmoko, Tatang Mitra Setia, Imran Razik, Carel P. van Schaik, Andrew Whiten, Paul-Christian Bürkner, Caroline Schuppli

## Abstract

Cultural learning permeates human skill and knowledge acquisition. To understand its significance for nonhuman animals, we must examine how social learning shapes the development of broad behavioral repertoires, and operates alongside other forms of learning. We analyzed nearly seven thousand records of behavioral indicators of social and individual learning across wild orangutans’ eight-year dependency period to assess their combined effects on ecological competence relevant for post-independence survival. Social learning was five times more frequent than individual learning and influenced subsequent learning for hours. Immatures varied substantially in their learning propensities and those with heightened social and individual learning propensities developed the most expansive diet profiles at independence. Our findings advance the science of cultural evolution by showing that multi-year social and individual learning synergistically shape great apes’ broad ecological competence.

---

Until as recently as the middle of the last century, culture was thought to be a feature of humanity that separates our species from the remainder of the animal kingdom (*1*). Recent research has revealed that to the contrary, culture defined as “information capable of affecting individuals’ behavior that they acquire from other members of their species through (…) forms of social transmission” (*2*), is widespread in the animal kingdom, extending to all main vertebrate taxa and even insects (*3–5*).

The evidence on which our understanding of animal culture is based, however, is very much a scientific patchwork based on a diversity of methodologies that have largely focused on short-term events in subjects’ lives and/or on relatively specific outcomes. Examples of research demonstrating animal culture based on short-term events in natural populations include studies tracing the diffusion of innovations across social networks, either spontaneous, as in the spread of moss-sponging in wild chimpanzees (*6*), or experimentally seeded, as in the case of foraging preferences in wild vervet monkeys (*7*). To demonstrate specific cultural outcomes in nature, a range of studies has shown that social learning contributes to the acquisition of preexisting complex behaviors, often using long-term, longitudinal datasets (*8–11*). Although longitudinal observational data are increasingly leveraged to investigate social learning in wild great apes, they have so far focused primarily on different types of tool use (*9*, *12–14*). Contrasting with these studies, two recent longitudinal studies in the wild have investigated possible social transmission of a broad competence – diet repertoire – but the evidence is either contextual (*15*) or drawn only from simulations (*16*). Beyond these studies on natural populations, there is a large corpus of rigorous experimental studies with captive populations, as in demonstrating the spread of challenging new tool use and other foraging techniques, from chimpanzees (*17*) to bumblebees (*18*). Numerous such studies now point to the existence of social learning capacities and pervasive cultural transmission in animals’ lives (*3–5*, *17–20*), but none capture the scale of what culture in the wild is thought to entail, in the round: a capacity to learn potentially very many things in the course of individuals’ development, generating a corresponding breadth of competence necessary for successful adult life.

Of course, animals likely also learn through their own efforts, such as targeted exploration and practice, without apparent immediate social influences, hereafter referred to as individual learning. While individual learning is thought to be less efficient and entail greater risks than social learning, such as poisoning and predation (*21*), a strict reliance on social learning can compromise individual’s adaptability to changing conditions (*22*) and may eventually lead to repertoire erosion due to incomplete transmission (*23*). Individual learning involves iterative problem-solving processes through which animals learn the affordances of their environment. By enabling direct assessment of current environmental conditions, individual learning enhances information reliability and provides more complete and up-to-date information about the payoff structure and variability of different behavioral options than socially transmitted information alone (*24*). Notably, social and individual learning may occur in tandem, making their distinction at times difficult (*15*). For example, animals may use the feedback generated by exploration and practice to extract additional information following social observation, particularly when learning and refining of complex skills (*19*). A handful of studies have accordingly begun to implement statistical models of “experience-weighted attraction” (EWA), which evaluate the relative roles of social and individual learning in the acquisition of novel tasks (*25–29*). The learning effects investigated in these studies were relatively short term, limited to the duration of the experiment, and to a single outcome of either tool use or the spread of a feeding technique.

To this day, our understanding of the combined effects of social learning and individual learning on individuals’ broad scale behavioral development remains minimal. Furthermore, whilst experimental studies provide evidence for individual differences in the use of social and individual learning (*9*, *30*, *31*), their findings are mixed as to whether an individual’s reliance on one form of learning predicts their reliance on the other (*32–34*). A negative correlation between social and individual learning propensities within individuals may arise from systematic differences among individuals in whether they predominantly pay attention to environmental or social cues (*33*), likely reflecting an underlying trade-off between investing in the two forms of learning. In contrast, a positive correlation may suggest that the same cognitive mechanism underlies the two forms of learning (*35*) and/or a stimulating developmental effect of social learning on individual learning abilities (*36*).

By linking individual variation in social and individual learning propensities to a behavioral outcome, we can evaluate whether there is a functional, adaptive trade-off or synergy between the two forms of learning. Accordingly, we have pursued a research programme that is uniquely expansive, in two main respects. First, we collected comprehensive measures of both individual and social information-seeking events of immature orangutans at different ages over the ∼8.5-year dependency period (fig. S1); and second, we evaluated the consequence of individual variation in the occurrence of these events for a global measure of “ecological competence” indexed by the breadth of dietary profile (number of identified, distinct food items eaten) achieved at independence (fig. S2 to S3). Adult orangutan diet encompasses a challenging total of around 250 different items selected from thousands of options in the forest (*16*), so our outcome measure of diet breadth is a significant indicator of the foraging competence that will determine how an individual fares post-independence, with consequent implications for its lifetime fitness.

We examined over 6,700 detailed all-occurrence records of peering behavior, as a measure of seeking information socially (Fig. 1A), and exploration behavior, as a measure of seeking information individually (Fig. 1B). These data were collected over 12 years of study and over 1,700 hours of detailed direct observation of 21 immature Sumatran orangutans (*Pongo abelii*) at the Suaq Balimbing research area, starting from their birth to the onset of independence. The dependency period ends at around 8.5 years of age when permanent association with the mother ceases and is the period during which most foraging competence develops (*37*). Peering is defined as sustained, attentive, close-range observation of the behaviors of conspecifics (*15*). The Suaq orangutans are estimated to peer for around 40,000 times over the course of their lifetime (*38*). That immatures follow peering by trying to repeat what they witness strongly suggests that individuals use peering to socially acquire ecological competence, including knowing what to eat and how to process the food items, sometimes involving tool use (*15*, *39*, *40*). We, therefore, used peering rate as a measure of an individual’s propensity to seek information socially, indexing social learning. Exploration is a key component of individual learning (*35*). We defined exploration as sustained—often destructive—manipulation of objects or attempted feeding without successful ingestion of food items (*41*). Age trajectories of the frequency, diversity, and persistence of exploratory object manipulation behaviors suggest that the orangutans at Suaq may also learn through such behavior (*41*, *42*). We thus used exploration rate as a measure of an individual’s propensity to seek information individually, indexing individual learning.

**Fig. 1.**
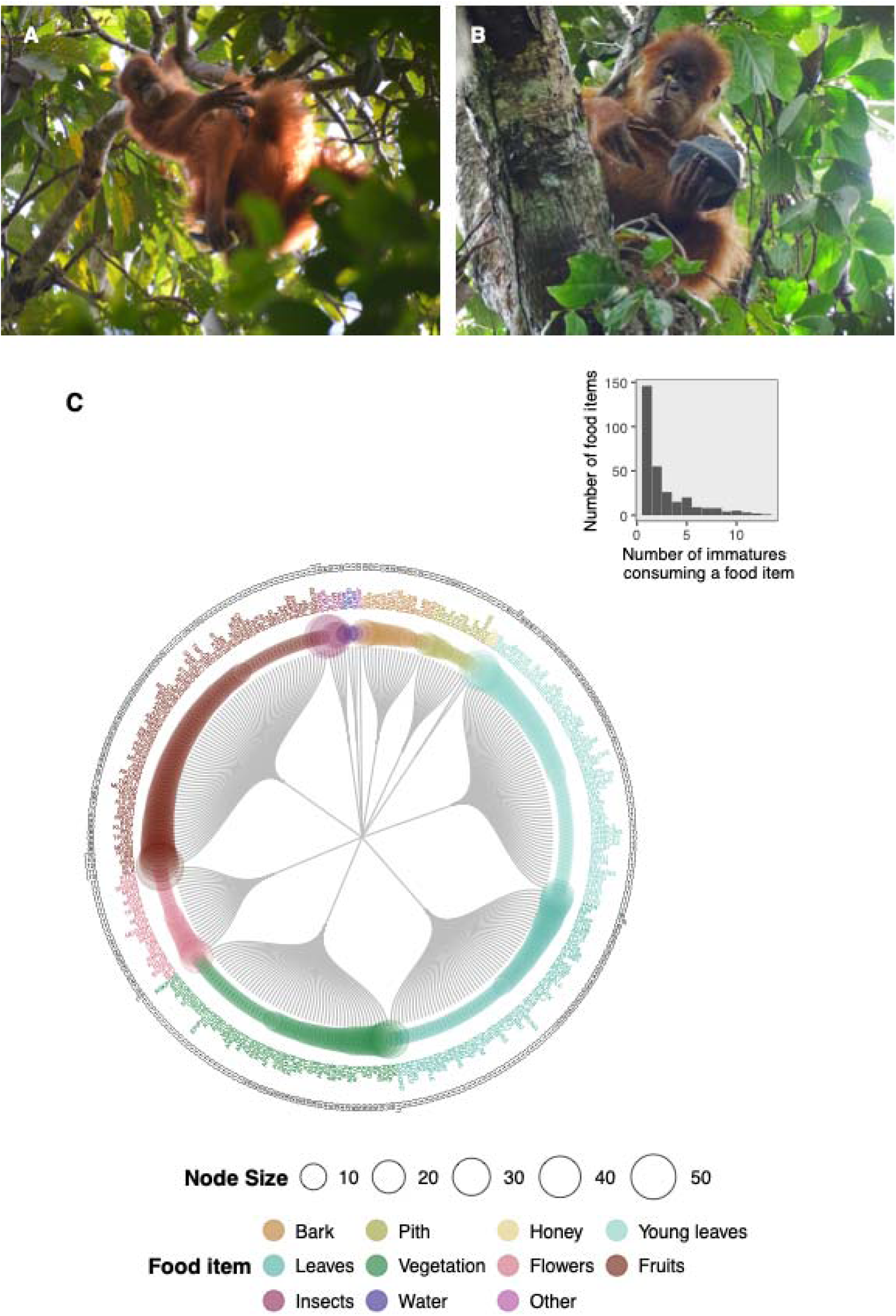
A Sumatran orangutan immature (**A**) *peering* at its mother, who is feeding on Neesia fruit and (**B**) *exploring* a Neesia fruit using a stick tool and (**C**) all the recorded unique food items (*N*=303, see table S1 for Latin names) eaten by a subset of 13 focal immatures for whom we had sufficient data for analyzing their ecological competence achieved at the onset of independence. Each node (colored circle) represents a food species. Each vertex (grey line) indicates a part of a food species. The node along with the vertex indicates a unique species-part combination (i.e., a food item). Note that multiple parts of a species were eaten by orangutans. Node size represents the recorded number of times each food item was consumed by these immatures during the study period. The number in the node label indicates how many (of the 13) focal immatures ate that specific food item. The origin (center) of the circular dendrogram does not carry any meaning in this figure. The inset shows the frequency distribution of the number of immatures consuming a food item across all food items (i.e., the extent of overlap in diet profiles among the immatures).

We hypothesized that immature orangutans engage in both individual and social forms of learning to develop broad ecological competence. Specifically, we tested whether:

i. Immatures engage more frequently in social or individual learning.
ii. Immatures show distinct individual propensities for both social and individual learning, and whether these propensities are correlated at the individual level.
iii. Immatureś levels of social and individual learning affect their development of ecological competence, and whether the effect of one learning form depends on the level of engagement in the other.

## Peering is more frequent than exploration and most exploration events are elicited by prior peering

Previous research suggests that peering elicits an immediate increase in exploration (*15*), which suggests that the two learning forms interact. We, therefore, conducted a hazard analysis to estimate the time point at which the instantaneous rate of exploration stabilized following a peering event, indicating a return to baseline exploration rates. We were able to incorporate an unprecedented number of peering (*N*=2113) and exploration (*N*=4649) events recorded across the 12 study years in our analysis (fig. S1). We used data only from immature individuals who had not yet reached independence from their mothers, meaning they were in constant association with them (*N*=21 immatures). On average, immatures peered at a rate of 1.25 events/h (SD=1.205 events/h) and explored at a rate of 3.71 events/h (SD=4.112 events/h), suggesting that they seemingly have a higher propensity to explore than peer. However, we found that most exploration events are far from truly individually initiated. We found that, in general, exploration events became independent of prior peering only at 145 minutes (fig. S4), highlighting a previously underexplored, intimate intertwining of social and individual information seeking at the immediate, proximate level. We refer to the exploration events that occurred within 145 minutes of the preceding peering event and before the subsequent peering event as socially initiated explorations (SIE; *N*=4235 events) and those that occurred at or beyond 145 minutes as individually initiated explorations (IIE; *N*=414 events). Immatures individually initiated explorations at a rate of only 0.25 events/h, (SD=0.801 events/h), which was one-fifth and one-sixth of the peering and SIE rates, respectively, indicating that IIE events are relatively rare in wild orangutans (fig. S5).

## Social and individual information-seeking propensities are individual-specific

We analyzed the total number of times immatures engaged in peering (*N*=260 focal follow days), SIE (*N*=257 focal follow days), and IIE (*N*=257 focal follow days) during each follow using three univariate Bayesian Generalized Linear Mixed effects Models (GLMM; U1, U2, and U3), controlling for immature’s age, sex, daily average number of association partners, monthly food availability, and observation duration, because these factors are known to have an effect on peering and exploration (*15*, *37*, *39*, *43*, *44*). We, therefore, refer to the resulting measures as the immatures’ *individual* propensities for peering, SIE, and IIE, which may arise from developmental and/or genetic influences. By including the number of association partners as a control in our model, we account for differences in social learning opportunities across the focal immatures. As these immatures live in highly overlapping home ranges (*45*), systematic habitat-induced differences in individual learning opportunities are not likely. We included a random intercept of immature identity to estimate social and individual information-seeking propensities for each immature and the variation in these propensities among immatures.

We found substantial variation among immatures in all three information-seeking propensities (table S2), suggesting that, on average, immatures differ from one another in their use of both social and individual information-seeking opportunities. Importantly, immatures differed from one another to a lesser extent in peering rate (SD of estimated individual average peering rate [95% CI]=0.69 [0.16, 1.42]; Fig. 2A) than in SIE rate (SD of estimated individual average SIE rate [95% CI]=1.24 [0.09, 2.80]; Fig. 2B) or IIE rate (SD of estimated individual average IIE rate [95% CI]=1.05 [0.11, 2.56]; Fig. 2C). When we performed sensitivity analyses using three new thresholds to classify exploration as independent of peering (70-minute, 130-minute, and 160-minute cut-offs; see Methods), the results showed that individual variation in IIE remained substantial but, with shorter time cut-offs, the variation became comparable to that observed in peering (see table S3).

**Fig. 2.**
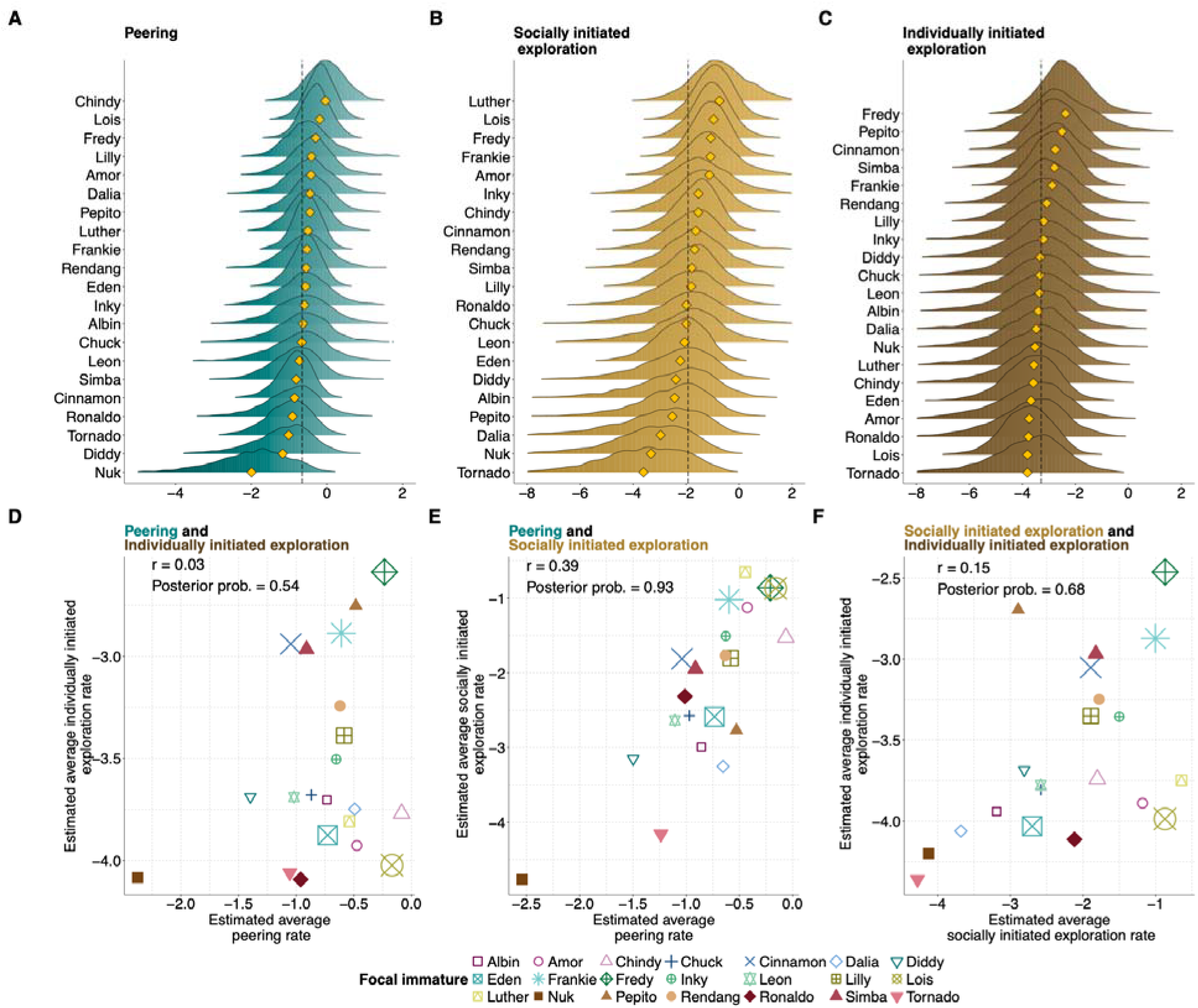
Substantial individual variation in information-seeking propensities and within-individual relationships between social and individual information-seeking propensities. Posterior distributions of the estimated average (**A**) peering, (**B**) socially initiated exploration, and (**C**) individually initiated exploration rates (per hour) for 21 immatures over their 8.5-year developmental period. The dashed black line represents the average over all immatures, and the yellow diamond represents the posterior mean of the estimate for each immature. Values to the left of the dashed line represent lower, and those to the right higher, than population average rates. Note that the x-axis scales are different among (**A**), (**B**), and (**C**). Scatterplots of individuals’ posterior mean for (**D**) peering and individually initiated exploration rates, (**E**) peering and socially initiated exploration rates, and (**F**) socially initiated and individually initiated exploration rates for the 21 immatures. Circle size represents the total observation duration for each immature between birth and 8.5 years of age (N = 2-360 hours). Correlation *r* and posterior probabilities for the hypothesis ‘*r* > 0’ are shown.

## Within individuals, information-seeking propensities are not correlated to each other

We fit three multivariate models to test for individual-level correlations among the following information seeking rates: peering events and IIE (M1), peering and SIE (M2), and SIE and IIE (M3). The models showed no evidence for a correlation between immatures’ different information-seeking rates (table S3; Fig. 2D-F). However, the posterior probability that the correlation between peering and SIE was greater than zero was 0.93, suggesting a positive association, although some uncertainty remains (Fig. 2E). Overall, these results suggest that immatures who peered at a higher rate did not necessarily socially or individually initiate exploration at a higher rate. However, a fourth multivariate model showed that peering and total exploration rates (the sum of SIE and IIE) were significantly positively correlated (M4; table S3). Ignoring the effect of peering on subsequent exploration can, therefore, erroneously suggest a positive correlation between rates of social and individual information seeking at the individual level. We then used the estimated individual peering and IIE rates from M1 as predictors of immatures’ ecological competence in our critical non-linear model to predict the effects of our measures of learning during development on ecological competence at the onset of independence.

## Inferring different forms of learning using long-term observations of wild individuals

We developed a new analytical approach to infer the contributions of different forms of learning to competence development in wild individuals, linking individual variation in information-seeking propensities during development to individual variation in broad-scale ecological competence at the onset of independence from the mother (Fig. 3).

**Fig. 3.**
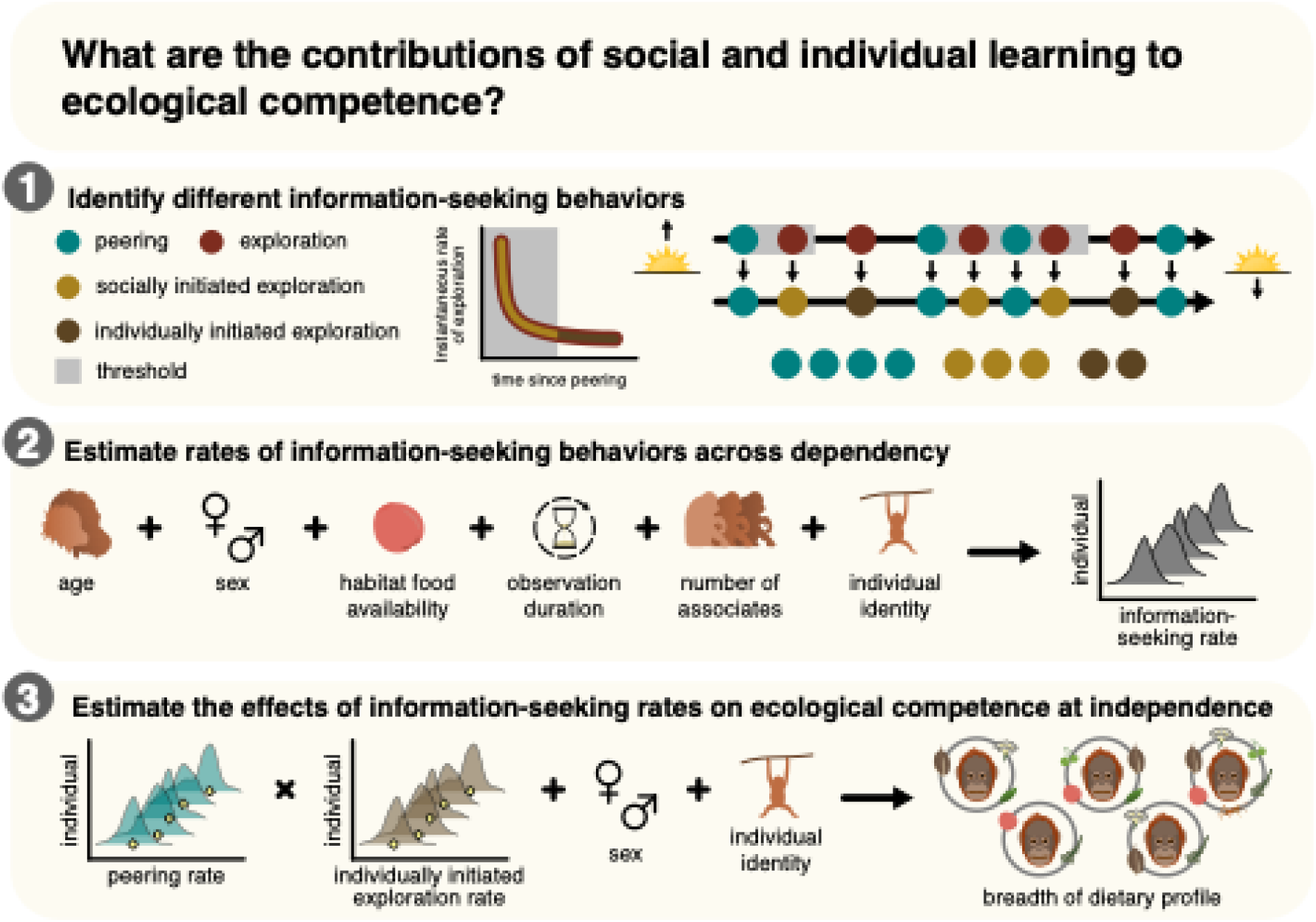
Testing the effects of social (peering) and individual (individually initiated exploration) information-seeking propensities on the development of ecological competence. Our approach was to first identify different information-seeking behaviors shown by the immatures, followed by the estimation of the average rates of these behaviors shown by each immature during development, to finally test the effects of these rates on the breadth of individual immature’s dietary profile at independence. In Step 1, the first row of dots illustrates immatures’ peering and exploration events, which were recorded by following individuals from their morning until the night nest whenever possible. The second row illustrates how the exploration events were classified as either socially or individually initiated exploration (SIE or IIE) events using a time threshold (grey). This procedure resulted in the identification of three different forms of information seeking, shown in the third row of dots. Step 2 illustrates the estimation of individuals’ information-seeking propensities across the dependency period, where we controlled for known confounding effects on information-seeking rates. Step 3 illustrates how we tested whether the breadth of the diet profiles at independence is predicted by individuals’ different information-seeking propensities, while controlling for sex as a known confounder of diet profile development. Steps 2 and 3 are depicted as statistical models, and the dependent variables of interest are to the right of the arrows.

Recognizing food items in their forest habitats is one of the most fundamental ecological competences wild orangutans need to develop to survive, especially when they become independent of their mothers and transition to a more semi-solitary lifestyle. Immature orangutans take ∼8 years (i.e., their entire dependency period) to reach the approximate breadths of the dietary profiles of their mothers (*37*). Because diet profiles are composed of distinct units (i.e., food items), their development is quantifiable. We thus used the breadth of an immature’s diet profile as a measure of its ecological competence. Overall, we identified a total of 303 unique food items eaten by 13 immatures during ∼4700 observation hours over 8.5 years of individual’s development prior to independence (Fig. 1C, fig. S2 to S3). Note that diet profiles could be assessed over more observation hours than peering and exploration rates, because only a subset of observers recorded all occurrences of the latter two behaviors. For eight of the 21 immatures that were used to quantify peering and exploration rates, we had too few observation hours to reliably calculate the age-dependent increase in the breadth of their diet profiles (see supplementary materials). Therefore, these eight immatures were not included in our subsequent analyses. By restricting our dataset to successful feeding events, rather than also including attempted feeding events, we investigated not only whether immatures learn what to eat but also how to process food to permit ingestion (*37*).

### Individual variation in learning propensities predict substantial differences in ecological competence at independence

We modelled the breadths of the dietary profiles that immatures achieved at independence as a function of their estimated average peering and IIE rates extracted from M1. Diet profiles broaden rapidly during early development and then plateau towards the end of the dependency period (fig. S3). We modelled this nonlinear effect of age through the Michaelis-Menten (MM) equation, implemented using a Bayesian framework. The breadth of the dietary profile at independence is represented by the *V*_max_ parameter of the MM equation, henceforth the parameter is called the breadth of the dietary profile. Based on our hypothesis, we modelled the breadth of the dietary profile as a function of the interaction between individuals’ estimated average peering and IIE rates (NM1; see Methods). When testing the effect of the interaction between peering and IIE on diet profile development, we may implicitly capture learning effects associated with SIE because of a putative positive association between peering and SIE rates (Fig. 2E).

The interaction between estimated average peering and IIE rates significantly predicted the breadth of the dietary profile at independence (table S4, Fig. 4A, where the effect size of the interaction effect is represented by the difference in the effect of peering on diet breadth across the three IIE levels). Specifically, we found a significant negative effect of the interaction on the breadth of the dietary profile at independence (Estimate [95%CI] = -36.65 [-70.11, -3.82] food items). Sensitivity analyses across different time thresholds to extract IIE (i.e., classify exploration as independent of peering; 70 minutes, 130 minutes, and 160 minutes) yielded consistent results (tables S5-S7), indicating robust findings.

**Fig. 4.**
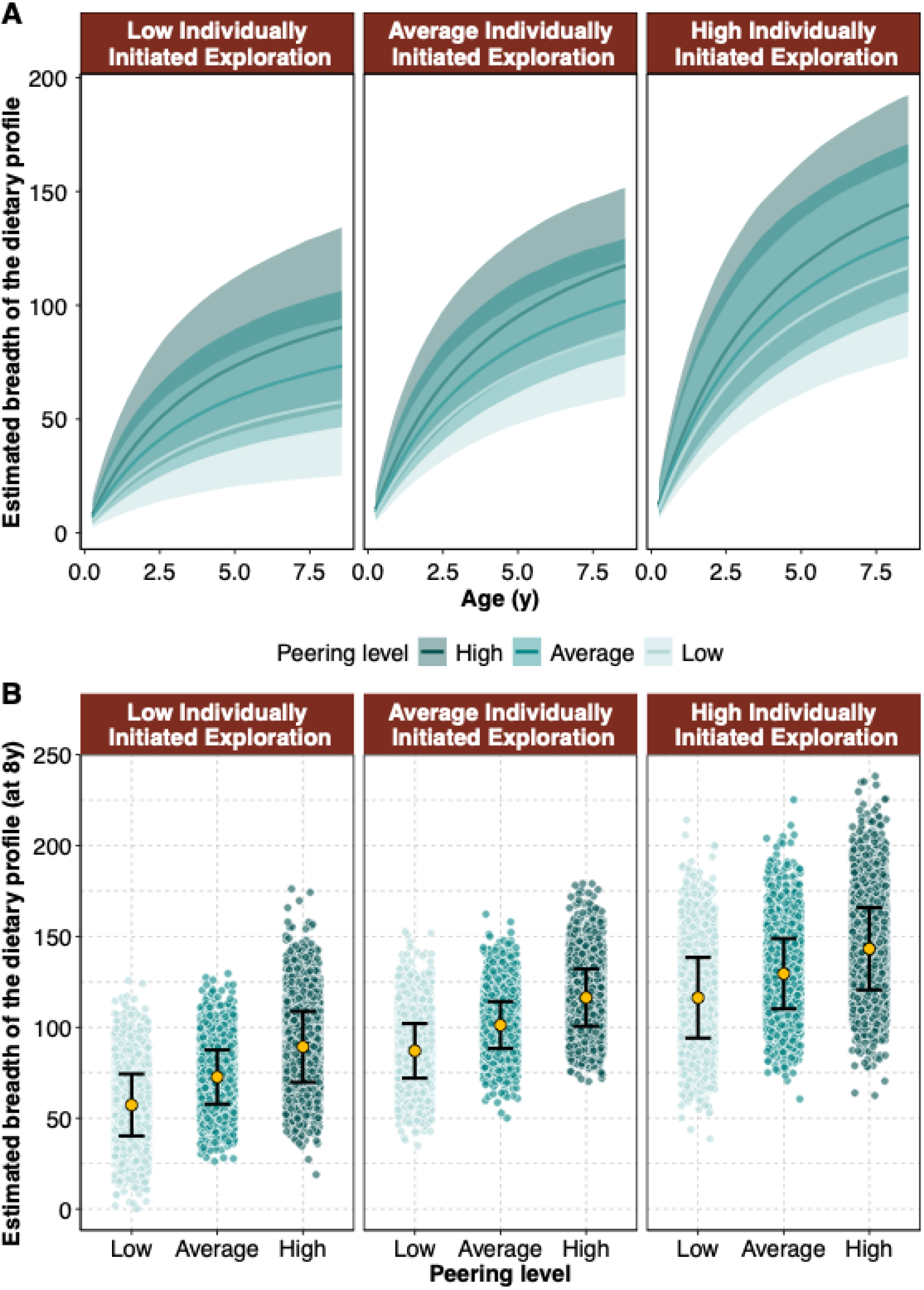
Effect of social and individual information-seeking propensities on the development of ecological competence. (**A**) Diet profile development of immatures between birth and onset of independence (model predictions from NN1 fit with Michaelis-Menten curves; breadth of the dietary profile was modelled as a function of a nonlinear interaction between peering and IIE; see Methods). (**B**) Estimated breadth of the dietary profile at independence as a function of individuals’ average estimated peering and individually initiated exploration rates. Low, Average, High refer to mean-1SD, mean, and mean+1SD. Each circle represents a posterior prediction. The yellow circle represents the posterior mean, and the vertical lines represent 95% CI.

### With increasing social and individual information-seeking propensities, immatures develop a broader ecological competence

Immatures with the highest peering and IIE rates developed the broadest diet profile by the onset of independence (Fig. 4A). In addition, the effect of peering on diet profile development became more pronounced when exploration rates decreased (negative interaction effect). Among immatures with the highest rates of IIE, those who peered at the highest rates had developed diet profiles that were 1.2 times the diet profiles of immatures who peered at lower rates (Fig. 4B). Under lower rates of IIE, the diet profiles of those who peered at the highest rates were 1.5 times larger than those who peered at lower rates (Fig. 4B).

To investigate the contribution of peering without follow up exploration to diet profile development, we first quantified peering events that were not followed by SIE. We found that only one-fourth (*N*=529) of all peering events were not followed by an SIE event. Furthermore, in only 15% of the follow days (*N*=39 days), a peering event occurred without any exploration event, showing that information seeking through peering alone is relatively rare (fig. S6). We also found that most of the peering events without a following SIE occurred towards the end of the dependency period (fig. S6). Finally, in our dataset we did not have any immatures that consistently peered without subsequent SIE (fig. S6). The rarity of peering events not followed by SIE prevented us from modelling the separate contributions of such peering to ecological competence development.

## Discussion

Ours is the first study of a wild nonhuman animal on the scale necessary to assess how episodes of both social and individual learning across the period of pre-independence shape the acquisition of a broad ecological competence. In orangutans, this spans an unusually long period in excess of eight years, at the end of which immatures begin to range independently of their mother. The expansive scale of our study yielded a series of discoveries that represent substantial new advances in the science of cultural evolution. First, we found that once we controlled for the potential confounding effects of age, sex, number of association partners, food availability and observation time, the remaining variation in what we called *individual* propensities for peering and exploration was nevertheless large. The relative expression of these different, major forms of learning thus appears to vary substantially between individuals, as has been shown to be the case in human children (*30*). Consistent individual biases in reliance on social learning have similarly been shown across repeated experimental tests in captive apes (*31*) and wild chimpanzees (*9*). Variation in individuals’ tendencies to engage in social versus individual learning, as well as in learning abilities, can arise from multiple factors, as suggested by studies across species. These include genetic differences (*46*), differential developmental influences such as early maternal care (*47*) and experienced differences in the reliability of social information (*48*), or a combination of genetic and developmental factors. In orangutans, where individuals primarily learn from their mothers (*15*) and where mothers systematically differ in their offspring-directed behaviors (*49*), differences in immatures’ learning propensities may be influenced by differences in maternal behavior. While studies on other taxa provide mixed evidence for covariation between individuals’ experimentally measured social and individual learning abilities (*32–34*), we found that individuals’ natural reliance on peering, IIE and SIE did not covary, although the covariation between peering and SIE needs further investigation. The absence of a negative correlation between individuals’ reliance on social and individual learning argues against trade-offs between investment in the two forms of learning and thus against strict behavioral specialization in the wild (*50*).

The substantial individual variation we documented highlights the importance of investigating (forms of) learning as individual-specific processes and not as traits that are expressed to the same extent across all individuals of a species. This should be taken into account whenever a study aims to compare different species’ tendencies for cultural or individual learning based only on averages. Individual variation in learning can emerge over evolutionary time if the benefits of learning are unpredictable, as suggested by a recent simulation study (*24*). We found that individual variation was lower in rates of peering than in IIE, suggesting there may a selection pressure on maintaining rates of peering, but not IIE, above a significant threshold. This may be because IIE likely carries significant risks in the wild (*21*). In addition, the higher individual variation in IIE rates compared to peering rates may partly stem from our choice of an empirically derived cut-off to classify exploration as independent of peering, as we found that the extent of individual variation in IIE, although substantial, decreased with shorter time cut-offs. Furthermore, we currently cannot rule out the possibility that variation in IIL partly reflects individual differences in the temporal structure of peering. We also found that individual variation was higher in rates of SIE than peering, which could stem from systematic differences in the complexity of the food items to which individual immatures are exposed. Previous research has found close similarities between mother-offspring diet profiles in wild orangutans, while mothers themselves differed in their diet compositions (*51*, *52*). Because orangutan diets vary in processing complexity, mothers may differ also in the overall complexity of their diet profiles. As less complex foods require less exploration after peering (*16*), immatures of such mothers might engage in lower levels of socially induced exploration (SIE), potentially contributing to greater variation in SIE than in peering. The level of individual variation in social and individual learning propensities observed in this study is surprising, as learning propensities are likely tied to fitness and, therefore may have important consequences in many species, including in non-WEIRD humans. Therefore, individual differences in learning merit further investigation, especially through longitudinal field studies, to understand their underlying causes, which may be environmental, developmental, and/or genetic (*53*).

A second major finding was that, in general, peering exerts an influence on exploration for as long as around two-and-a-half hours. This leads us to suggest a distinction between what we label here as “Socially Initiated Learning” (SIL) stimulated by peering within this period—which we accordingly place within the broader conception of social learning (Fig. 5)— and “Individually Initiated Learning” (IIL) that occurs at other times. In Sumatran orangutans, a dependent immature spends more than 50% of its time within 10 m of its mother until at least six years of age (*54*). A previous study on peering and exploration in this population found that the rate of exploratory behavior directed at a food item was higher in the hour after, compared to before, peering at one’s mother consuming the same food item (*15*). Our results suggest that peering may direct an immature’s attention to specific locations and/or items. This, in turn, indicates that SIL is consistent with observational social learning or indirect learning processes such as local or stimulus enhancement (*19*). The important distinction between SIL and IIL should now be taken into account when refining the “Experience-Weighted Attraction” models that are beginning to proliferate. Hitherto they have typically made only a simple binary distinction between events labelled as social versus individual (or truly asocial) (*25–29*), whereas we find that the greatest proportion of exploration, in the form of SIL, is consequent on peering and thus falls at the intersection between social and individual learning. Correlations between social and individual learning at the individual or population level must be thus interpreted with caution to avoid false positives. SIL occurs at a rate six times higher than IIL and likely involves selective exploration of the peered-at-food-item (*15*). It is easy to appreciate that learning through peering alone, what we call Strictly Social Learning (SSL), and SIL may act synergistically in diet profile development. Peering stimulates SIL, and through effects such as prompting individuals to practice recently observed activities, allows developing individuals to gradually perfect their skills, including complex skills such as tool use (*55*). Indeed, our study demonstrates that during most of the developmental period, immatures engage in both SSL and SIL, which poses a challenge in estimating their individual effects. However, that these two forms of learning often co-occur already provides evidence that both are important for the development of ecological competence in Sumatran orangutans.

**Fig. 5.**
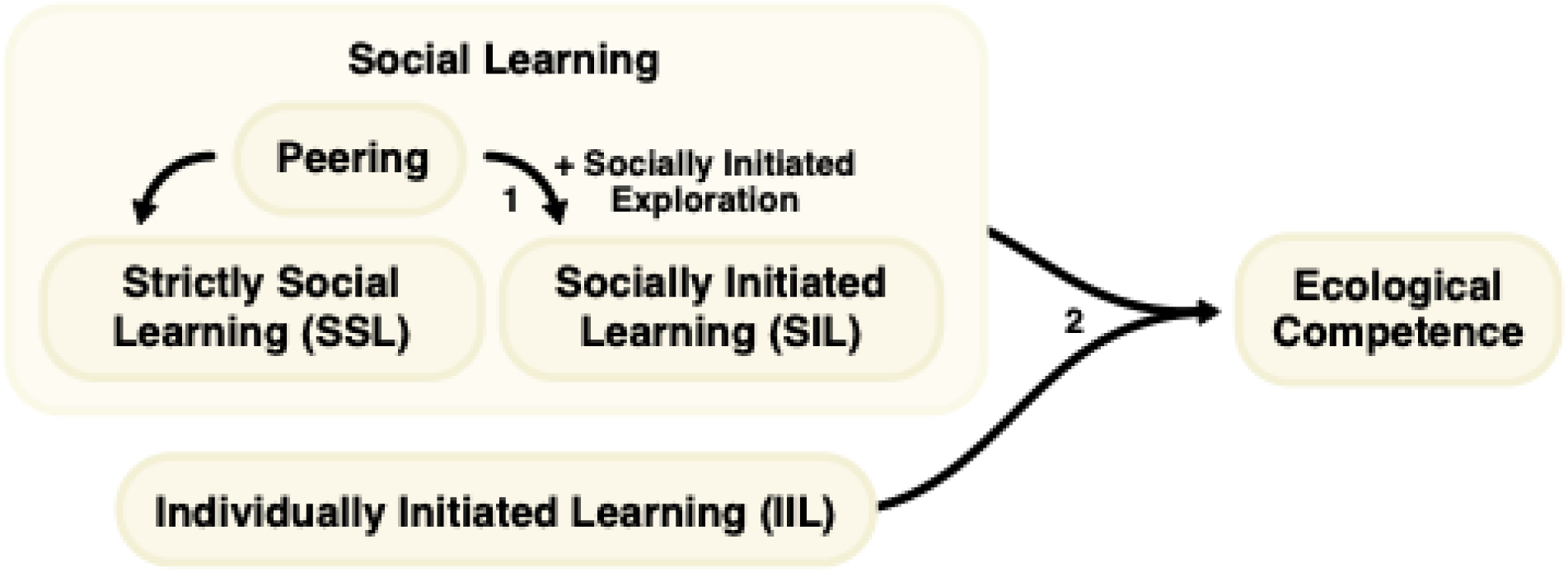
Overview of the forms of learning involved in diet profile development in Sumatran orangutans. 1. Peering triggers socially initiated learning and 2. Social learning and individually initiated learning synergistically influence ecological competence development.

Our third, overarching finding is that variation in the totality of immatures’ social and individual learning shapes the ecological competence that is so critical for achievement of independence. We believe this constitutes the most important contribution to the field of cultural evolution. Whilst there are other competencies a young ape must learn, acquiring ecological competence is likely directly relevant for lifetime fitness and includes the learning of a broad range of skills and knowledge types. We found that ecological competence, indexed by the breadth of the dietary profile, is predicted by an interaction between observational social learning through peering (which includes SIL and SSL) and IIL. At all levels of IIL, increases in social learning through peering led to greater ecological competence but the effect of peering on ecological competence became more pronounced as IIL decreased. This suggests that social learning through peering and IIL work synergistically in the development of diet profile (Fig. 5). Moreover, when immatures do not engage in heightened individual learning to acquire a competence, social learning through peering may compensate for the lack of individual effort.

In contrast to SSL and SIL, cases of IIL are distinguished by their lack of direct behavioral linkage to peering, but we suggest three potential answers for how and why IIL interacts with peering to enhance ecological competence. At the developmental level, increased engagement in social learning expands an individual’s pool of affordances, thereby enhancing their overall learning ability, as predicted by the Cultural Intelligence Hypothesis (*36*, *56*). In our study, immatures with increased social learning through peering and IIL rates developed the broadest diet profile. However, we did not find a positive correlation of peering and IIL within individuals, which would be expected if these developmental effects were driving the interaction between IIL and peering. Part of the answer may be that memory of earlier peering still shapes their content well beyond the empirically derived cutoff, which may be why we find a positive correlation between peering and total exploration (sum of SIE and IIE). If this is true, our estimate of the relative importance of social versus individual learning is likely a lower bound, and the influence of social learning on the development of ecological competence may be even greater than what is currently estimated. In addition, IIL rates may reflect individual differences in behavioral persistence or sustained engagement with objects during periods without recent peering.

More broadly, individual differences in activity levels or general learning ability may underlie engagement in both social learning and IIL, although not necessarily to the same extent, potentially resulting in a synergy between the two. A third explanation for the interaction between peering and IIL may be that the temporal gap may facilitate adaptive exploration of behavioral variations at the fringes of what had been learned through peering. Truskanov and Prat recently suggested that in an uncertain world, copying observed actions imperfectly, coupled with a degree of trial-and-error learning, may be the most adaptive strategy (*22*). Their simulations support this idea and highlight trial-and-error learning as a stabilizing force in cultural transmission. This is likely relevant to our orangutan population and indeed to many other species.

The cultures of the great apes have been shown to be the most complex among nonhuman animals in the extensive arrays of traditions that differ between communities (*57–59*). Research on chimpanzees led these studies in the later part of the twentieth century (*60*) but the more recent recognition and validation of peering as an index of social learning in orangutans (*15*), much reinforced by the present study, indicates that culture pervades apes’ lives to a greater extent than the earlier studies had indicated, with individuals of the Suaq population learning socially in as many as 195 different contexts (*38*). The present study extends such understanding of the pervasiveness of cultural learning in great apes into another dimension, documenting episodes of peering across eight years of immatures’ development and establishing their impact on achieving the extensive and crucial dietary breadth that sustains their mature, independent foraging. Other, more focused studies have shown that cultural learning additionally pervades apes’ later lives too, as when maturing males selectively peer at adult males, the optimal models for their larger-bodied mature foraging needs (*39*), and at resident (“expert”) individuals when they migrate into new ranges (*40*). Importantly, the study also highlights that cultural learning acts in synergy with individual learning.

Our overall findings show that different forms of learning act synergistically to shape great apes’ behavioral repertoires, with greater social learning consistently associated with larger repertoires across all levels of individual learning. Our study can thus be summarized as demonstrating the pervasiveness of the social effects on learning in great apes’ lives and also a more nuanced reality in terms of the linkages between the social and individual components of learning. All of this brings great ape cultural learning mechanisms closer to the complexities of the underlying mechanisms of human culture. Since orangutans, the other great apes and humans share a common ancestry, it is apparent that the extraordinary heights of human cultural learning were likely built on already substantial evolutionary foundations.

## Acknowledgments

We thank all the dedicated field staff, students, and research assistants – in particular, Anais van Cauwenberghe, Ellen Meulman, Lara Nellissen, Natalie Oliver Caldwel, Sofia Forss, – who, besides CS, collected most of the all occurrence data on peering and exploration used in this study. We are grateful to the Indonesian State Ministry for Research and Technology (BRIN-RISTEK), the Directorate General of Natural Resources and Ecosystem Conservation–Ministry of Environment and Forestry of Indonesia (KSDAEKLHK), the Ministry of Internal Affairs, Indonesia, the Sumatran orangutan conservation Program (SOCP), Balai Besar Taman Nasional Gunung Leuser (TNGL), in particular Arif Saifudin, Zakir, and Samsul Amar, and all staff members in Medan for their permission and support to conduct this study. We thank Universitas Nasional (UNAS) for their support and collaboration. We thank Elliot Howard-Spink for useful discussions about the analysis. We thank Tri Rahmaeti for coordinating our research permits and liaising with Indonesian offices and institutions. We thank the technical assistants, Richard Young and Francois Lamarque, for maintaining the SUAQ database.

## Funding

Marie Sklodowska-Curie Actions postdoctoral fellowship (TR)

Max Planck Institute of Animal Behavior (CS)

University of Zurich (CS)

A.H. Schultz Foundation (CS)

Leakey Foundation (Primate Research Fund and project grant) (CS)

SUAQ Foundation (CS)

Volkswagen Stiftung, Freigeist fellowship (CS)

Stiftung für Mensch und Tier, Freiburg i.Br. (CS)

## Author contributions

Conceptualization: TR, CS

Data collection: CS

Methodology: TR, PB, CS

Data curation: TR, MTW

Formal analysis: TR

Validation: TR, PB

Investigation: CS

Visualization: TR, IR

Funding acquisition: TR, CS

Project administration: SSUA, TMS, CS

Supervision: PB, CS

Writing – original draft: TR, AW, CS

Writing – review & editing: TR, MTW, SSU, TMS, IR, CPvS, AW, PB, CS

## Data and materials availability

All data and codes are available are available on Dryad data repository (*61*).

## Materials and Methods

### Data Collection

This research was approved by the Indonesian State Ministry for Research, Technology and Higher Education, the National Research and Innovation Agency (BRIN RISTEK; Research Permit No.: 152/SIP/FRP/SM/V/2012), the Directorate General of Natural Resources and Ecosystem Conservation under the Ministry of Environment and Forestry of Indonesia (KSDAE-KLHK) and the Gunung Leuser National Park (TNGL). This research was conducted in accordance with the Indonesian law and the ethical standards for research on non-human primates, following the Guidelines for the Treatment of Animals in Behavioural Research and Teaching of the Association for the Study of Animal Behaviour (*62*).

We investigated whether immatures I. engage more frequently in social or individual learning, II. show distinct individual propensities for both social and individual learning and whether these propensities are correlated at the individual level, and III. whether immatures’ levels of social and individual learning affect their development of ecological competence, and whether the effect of one form of learning depends on the level of engagement in the other in Sumatran orangutans (*Pongo abelii*). To address these questions, we used longitudinal and cross-sectional data collected from 2007 to 2019 at the Suaq Balimbing research area in the Gunung Leuser National Park (3°42 N, 97°26 E) in South Aceh, Indonesia. As this study was observational in nature and relied on identifying individual orangutans in the field, blinding to individual identity was not feasible.

CS and a total of 15 other observers collected data on daily behavior through instantaneous scan sampling at 2-minute intervals during focal animal follows, following a standardized protocol for orangutan data collection (www.ab.mpg.de/571325/standarddatacollectionrules_suaq_detailed_jan204.pdf). To quantify social and individual information-seeking propensities, we included *all occurrence* data on peering and exploration, which was collected by a subset of observers. Focal individuals were followed from the morning nest (∼06:00) until the night nest (∼18:00) when possible. When a focal individual could not be followed nest to nest, the follow type fell into any of the three following categories: found-to-nest, found-to-lost or nest-to-lost. All the researchers underwent a training period, and the data collected by a new researcher was included for analysis only when the inter-observer reliability reached a minimum of 85% with experienced researchers for simultaneous focal follows. In addition, we controlled for potential observer-to-observer differences in our models (see statistical analyses). Data collectors were blind to the aims and objectives of the current study.

We used peering as a behavioral indicator of social information seeking. Previous research on the study population showed that peering occurs most frequently in learning-intense contexts, such as feeding and nesting (*15, 63*). We defined <u>peering</u> as attentive and sustained (≥5 seconds) close-range (2-5 m) observation of conspecifics’ behavior (*15*). We recorded a total of 2113 peering events (across all behavioral contexts) by 21 focal immatures (*N*=5 females and 16 males; note that the sex-ratio among the immatures was naturally skewed in the study area at the time of data collection) between 0.51 and 8.72 years of age (fig. S1) during 1759.98 *h* of observation (*N*=260 follows). On three of the observation days, data on peering but not on exploration were collected (see below; the observer did not reach sufficient inter-observer reliability for exploration yet). Since our main aim was to assess individual’s overall propensities to peer and explore, slight differences in the exact dates of data collection are unlikely to affect our results.

We used exploration as a behavioral indicator of individual information seeking. We defined <u>exploration</u> as sustained manipulation of objects or attempted feeding without successful ingestion of food items (*15, 41*). Object explorations were often destructive in nature. We recorded a total of 6125 exploration events by 21 focal immatures between 0.51 and 8.72 years of age (fig. S1) during 1743.2 *h* of observation (*N*=257 follows). Through exploration, individuals can find out whether an object is edible or not or contains a food item, i.e., whether an object is indeed a food item or a substrate of a food item (e.g., dead wood or tree hole, which often contains insects; or the pods of a pitcher plant, which contain water). Furthermore, individuals may explore items such as sticks, which may help them learn how to use these objects as tools to access particular food items. Note that as we analyzed all exploration events and not only the events the involved the same contents that were peered at, the average estimated rates (see statistical analyses) represent overall propensities to explore rather than content-specific performance or practice of behavior.

All but two of the 21 immatures survived to weaning age. One dependent immature (Rendang) died following his mother’s death at the age of three years. The other immature (Leon) died within one year of birth. He failed to thrive; however, his peering and exploration rates were close to population average and not very low when compared to immatures who survived the entire developmental period (see Main text, Fig. 2). These two immatures, along with six others (eight immatures in total, see below), could not be included for the subsequent analyses on ecological competence development due to sample size limitations.

Adult Sumatran orangutans at Suaq have large diet profiles, including ∼250 unique food items (*16*). At each feeding activity *scan*, we noted down the species and the parts eaten by the focal individual. Following previous research, we divided parts of the species eaten into broad categories of flowers, fruits, young leaves, mature leaves, vegetation, bark, pith, honey and insects (*37*). We considered each ingested species-part combination as one unique <u>food item</u> (*37*). Note that we distinguished between young leaves and mature leaves (Fig. 1C). Leaf type can be visually recognized by the shade, with young leaves usually being lighter in color than mature leaves. We distinguished between the two because adult orangutans in this population show a clear preference for the former over the latter for most plant species: of the 38,771 feeding events on leaves, 28,515 of them (or 73.5%) were on young leaves. Vegetation (Fig. 1C) is inclusive of all vegetative matter that does not fall into any of the other categories (e.g., the stems of vines). Therefore, our count of the number of unique items eaten by an immature is a conservative estimate and does not lead to any overlap between any of the categories of food items. We identified a total of 303 unique food items eaten by 13 focal immatures between 0.26 and 8.57 years of age (fig. S1) during 4792.43 *h* of observation (*N*=453 follows). We only considered feeding scans during which the immatures properly ingested the food items without the help of their mothers. We recorded a total of 51,055 feeding scans, out of which the food item was identified in 50,542 scans resulting in 1% of the feeding scans with unidentified food items (across 453 follows; fig. S2). Note that diet profiles could be assessed over more observation hours than peering and exploration rates, because only a subset of observers recorded all occurrences of the latter two behaviors.

Each of these 13 immatures was observed during at least 2 different age years (range: 2 – 8.5 observation years), and each focal follow was at least 5 *h* long. When we adjusted this cut-off to be greater than 5 hours, an increase in follow hours did not result in an increase in the recorded number of unique food items eaten by a focal immature. Using all the feeding scans with an identified food item, we calculated the breadth of the dietary profile as the cumulative number of unique food items eaten by each immature at each sampled age (fig. S3), which was our measure of <u>ecological competence</u>. Food species were identified by the observers with the help of long-term local field assistants, all of whom had multi-year experience in identifying food species at the research site. To assess inter-observer reliability in food species recognition, we compared feeding events from 40 simultaneous follows (recorded during the study period), during which no communication took place between the observers, as these follows were part of interobserver reliability tests. These follows included 4168 feeding events, and the observers independently arrived at the same species-part combination for 3696 of these events. This leads to an inter-observer reliability of 87.3% for food item identification during these follows. Note that these tests were conducted during the end of the training periods of new observers and thus represent food item recognition skills of new rather than experienced observers. For eight of the 21 immatures that were included in our analyses on peering and exploration rates, we had too few observation hours (i.e., the lengths of the focal follows were <5 h and/or the immatures were followed during only one sampling year) to reliably calculate the age-dependent increase in the breadth of their diet profiles. Therefore, these eight immatures were not included in our final nonlinear model to assess ecological competence.

Within the home ranges of frequently observed focal individuals, we monitored all the trees with a DBH (Diameter at Breast Height) of >10 cm along two long-term phenology transects (North-South and East-West) for the presence of unripe, semi-ripe and ripe fruits. Each month, we quantified the <u>Food Availability Index</u> (FAI) from these ∼1000 trees as the percentage of fruiting trees in these phenology plots (*64*).

We calculated daily average association size for each follow by taking the mean of the number of association partners (defined as individuals within 50 m of the focal individual) across all the scans taken during a follow.

### Statistical analyses

All the statistical analyses and plotting were performed using R, version 4.3.1 (*65*). All the graphs were generated using the *ggplot2* package (*66*) (version 3.5.2) or *ggraph* package (*67*) (version 2.2.1). In our Bayesian generalized linear mixed-effects models (GLMM), we considered an effect to have strong evidence for being non-zero when its 95% credible interval did not include zero.

### When does an exploration event become independent of a peering event?

From previous research on the study population, we know that peering triggers increased selective exploration in immatures (*15*). To determine the timepoint after which exploration rates return to baseline following a peering event, we first calculated the time to each exploration event following a peering event and before the next peering event. We then estimated the hazard function of exploration events over time since the last peering event. To do so, we used the *muhaz* package (*68*) (version 1.2.6.4) in R, which implements kernel-smoothed hazard estimation for time-to-event data.

To identify the point at which exploration rates (i.e., the “hazard”) stabilized—signifying independence of exploration events from the preceding peering event—we computed the first derivative of the hazard function (i.e., its slope) across time. A sustained near-zero slope indicates a flattening of the hazard curve, meaning that after this point in time the probability of exploration is no longer dependent on the last peering event. To avoid spurious identifications due to short-term fluctuations in exploration rate, we checked whether the near-zero slope remained stable for at least 25 continuous timepoints. We implemented this condition using a moving window approach using the *zoo* package (*69*) (version 1.8-14). Based on the estimated time points, we split the exploration events into two: socially initiated exploration events (SIE, ≤145 minutes since the preceding peering event and till the next peering event) and individually initiated exploration events (IIE, >145 minutes since the preceding peering event and till the next peering event).

Of the follows used for the analysis, 77% were nest-to-nest follows. For follows that did not begin at an immatures’ morning nests, we excluded exploration events that occurred before the first peering event of the day from the calculations of exploration events independent of peering. In addition, since not all follows ended at immatures’ night nests, to avoid any systematic variation across the follows or immatures, we calculated the number of exploration events that were independent of peering using only the exploration events that occurred between two peering events, regardless of the follow type (*N*=4649 exploration events). This did not affect our sample size (i.e., focal follow days).

All the models were fit using a Bayesian approach using *Stan*, version 2.32.2 (*70, 71*) through the *brms* package, version 2.22.0 (*72*). All the models were run for 4 chains with 2000 iterations each, out of which 1000 were warm-up iterations, leading to 4000 samples for each model estimate. To prevent divergent transitions, we set adapt_delta to 0.99, step size to 0.01 and maximum tree depth to 15. We used the ‘bayes_R2’ function of the *brms* package to calculate *R*^2^ to assess model fit (*73*).

### Predictions I & II Do immatures engage more frequently in social or individual learning and show distinct individual social and individual learning propensities that are correlated at the individual level?

We first fit four univariate Generalized Linear Mixed Effects Models (GLMM), for peering (U1), SIE (U2), IIE (U3), and total exploration (U4), with all the fixed and random effects (see below) to find out whether there was variation amongst immatures in their information-seeking propensities. We further performed sensitivity analyses (see below). The number of recorded information-seeking events during each follow was the dependent variable in each of the models. We refer to these as full models. We then fit reduced models, which lacked the random effects of immature ID but contained all other fixed and random effects. We refer to these as reduced models. We compared the full and the reduced model using leave-one-out cross-validation (*74*) through the *loo* function of the *brms* package and considered individual variation in information-seeking propensities to be substantial if the full model had the highest ELPD value – i.e., the highest predictive value.

Since there was substantial variation among immatures in all four information-seeking rates, to determine whether peering and exploration rates were correlated at the immature-level, we fit multivariate GLMMs. Here again we used the number of recorded information-seeking events during each follow as the dependent variable. We fit four multivariate models based on datasets including: M1. peering and IIE events, M2. peering and SIE events, M3. SIE and IIE events, and M4. peering and total exploration events.

We fit the models using a negative binomial error distribution and a log link function. The multivariate model allowed us to test whether there was a correlation between different *estimated average rates* while accounting for the error associated with these estimates. To this end, we did a directional hypothesis test to check whether the correlation estimate (*N*=4000 samples) was different from 0 (*r*>0) using the ‘hypothesis’ function of the *brms* package. If the correlation estimate was greater than 0, we inferred that information-seeking propensities were correlated at the immature level. We also performed three sensitivity analyses (see below).

The fixed and random effects included in the univariate and multivariate models are detailed below.

#### Fixed effects

Since the observation duration varied across the follows, we used it as an offset, supplied through the ‘rate’ function of the *brms* package. We included age and sex (male/female) of the focal individual, FAI, and daily average association size during a follow as predictors of information-seeking rates to control for their influence on these variables (*15, 37, 39, 43, 44*). Since peering peaks around 3-4 years of age in the study population (*15*), we additionally included a quadratic effect of age in the peering model. As there were slight differences in the number of follows used for peering *vs* exploration (N=260 *vs* 257), we used the ‘subset’ function of the *brms* package to choose the appropriate follows for analyzing each behavior. For easier interpretation of model estimates and to ease model convergence, we *z*-standardized the continuous predictors (age, daily average association size, and FAI) to a mean of 0 and a standard deviation of 1 and dummy coded and centered the categorical predictor.

#### Random effects

We used maximal random effects structure in all the univariate and multivariate models. We included immature identity as a random intercept, as we were interested in the random intercept estimates, and age, daily average association size and FAI as random slopes within immature identity. The random intercepts from the peering model represent the *estimated average peering rates* (i.e., overall propensity to peer) and those from the exploration models represent the *estimated average SIE, IIE, and total exploration rates* (i.e., overall propensity to explore) for each focal immature during development. As we included *all occurrence* data, we included observer as an additional random intercept to control for observer-to-observer differences in sampling.

#### Priors

We used the default flat prior in *brms* for the univariate and multivariate models to avoid imposing unwarranted assumptions on the model.

We assessed model convergence through R-hat values (*75*), which were equal to 1.00, and bulk and tail ESS, which were all >1000. ESS refers to Effective Sample Size, which is a measure of how much independent information there is in autocorrelated chains. It indicates how well the algorithm explores the posterior distribution, and values higher than 200 indicate better mixing and more reliable parameter estimates. We also visually inspected the trace plots and found no model convergence issues.

We used the estimates from M1, based on the 145-minute cut-off dataset, for the subsequent nonlinear analyses (NM1, NN1; outlined under III). We present and interpret the results of these analyses in the main text.

### Prediction III Do the levels of social and individual learning affect the development of ecological competence and do the effect of one learning mode depends on the level of engagement in the other?

To investigate the development of ecological competence, we analyzed how the breadth of the diet profile develops in immatures. We modelled diet profile size (i.e., the cumulative number of unique food items eaten by an immature during the developmental period) as a nonlinear function of age using the Michaelis-Menten (MM) equation (*76*). The MM equation is used in enzymatic kinetics to describe the transformation of a substrate into a product with the help of two parameters, *V*_max_ and *K* [1]. The curve described by the MM equation is a rectangular hyperbola that rises steeply at the beginning and gradually levels off toward a maximum. Previous research shows that in immature orangutans, the breadth of the diet profile follows this trajectory, increasing with age but approaching an upper asymptote towards independence (*30*).

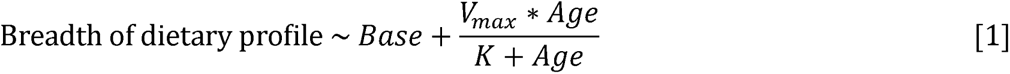

#### Predictors

In the context of diet profile, *V*_max_ represents the maximum breadth of the diet profile or the cumulative number of unique food items that an immature has eaten by the onset of independence (∼8.5 years in the current dataset). This is the upper asymptote of the MM curve. *K* represents the half-saturation age or the age at which an immature reaches 50% of *V*_max_. *K* controls how quickly the breadth of the diet profile approaches *V*_max_. We modelled *V*_max_ and *K* with a softplus transformation to constrain them to positive values. We modelled *V*_max_ as a function of an interaction between estimated average peering rate and estimated average IIE rate during development (obtained from M1) to predict the interactive effect of social and individual information-seeking rates on ecological competence. We scaled the estimates using the ‘scale’ function to ease the interpretation of model coefficients. We further performed three sensitivity analyses (see below).

Since previous research found that the breadth of the diet profiles differed between male and female immatures (*52*) we modelled both *V*_max_ and *K* as a function of immature sex. As there were repeated samples for each immature, we included the random intercepts of immature identity in these two parameters. We refer to this model as NM1. Apart from this model with a linear interaction term, we built another nonlinear model (NN1) where we modeled *V*_max_ as a nonlinear function of these rates, using a smooth interaction term s(peering, individually initiated exploration, k = 5), where k specifies the number of basis functions used to approximate the smooth surface (*77*). This provides a balance between model flexibility and parsimony, allowing moderate curvature while reducing the risk of overfitting. In this way, the nonlinear interactions allowed us to capture the complex, interdependent relationship between social and individual information-seeking rates on diet acquisition, which are unlikely to interact in a strictly linear fashion. For example, the benefit of peering may depend on the level of exploration, or *vice versa*, which is impossible to be captured by a simple, linear interaction term. Since the estimates of the model with the smooth interaction term is not interpretable, we report the model estimates from NM1 but present the graph based on NN1.

#### Random effects

We included immature ID as a random effect for the *V*_max_ and *K* parameters.

#### Priors

We used weakly informative priors to regularize parameter estimation while allowing the data to dominate. For the baseline diet breath (base), we set a truncated normal prior of N(0,5), as immatures are dependent on their mothers for nutrition at birth. We set normal priors of N(150,10) for the intercept, as the maximum cumulative diet breadth was ∼150 in our data set (fig. S3) and previous simulation-based research found that immatures’ diet breadth reaches up to ∼225 (*16*); N(15,4) for the effect of sex, as previous research found a 10% difference in diet breadth between the sexes towards independence (*52*); N(0,4) for estimated peering and IIE rates; and N(0,10) and N(0,2) for the random intercept of immature identity for *V*_max_ and *K*, respectively. We set normal priors of N(2,1) and N(1,0.5) for the intercept and the effect of sex, respectively, for the *K* parameter, as it can be discerned from previous research that immatures reached 50% adult-like diet breadth between 2 and 4 years with a 1-2 year difference between males and females (*37,52*). We selected these priors to reflect biologically plausible ranges while minimizing undue influence on the posterior. We did not set informative priors for the interaction between peering and IIE because no prior knowledge of this effect was available. Instead, we used the default flat prior in *brms* for the interaction term to avoid imposing unwarranted assumptions on the model.

All R-hat values were equal to 1.00, and all bulk and tail ESS were >2000. Visual inspection of the trace plots showed no model convergence issues.

### Sensitivity analysis

To ascertain that our results are robust and that they are not an artefact of the specific time cut-off of 145 minutes (used to classify exploration as independent of peering), we performed sensitivity analysis using three other time cut-offs: 130 minutes (15 minutes before), 160 minutes (15 minutes after), and 70 minutes (obtained by relaxing the number of timepoints for which the slope of instantaneous exploration rate against time since peering should be near-zero; relaxed from 25 (see above) to 10 timepoints). With these cut-offs, we ran additional sets of univariate models (SU1, SU2, SU3), multivariate models (SM1, SM2, SM3), and nonlinear models with a linear interaction between peering and IIE rates (SNM1, SNM2, SNM3).

---

**Fig. S1.**
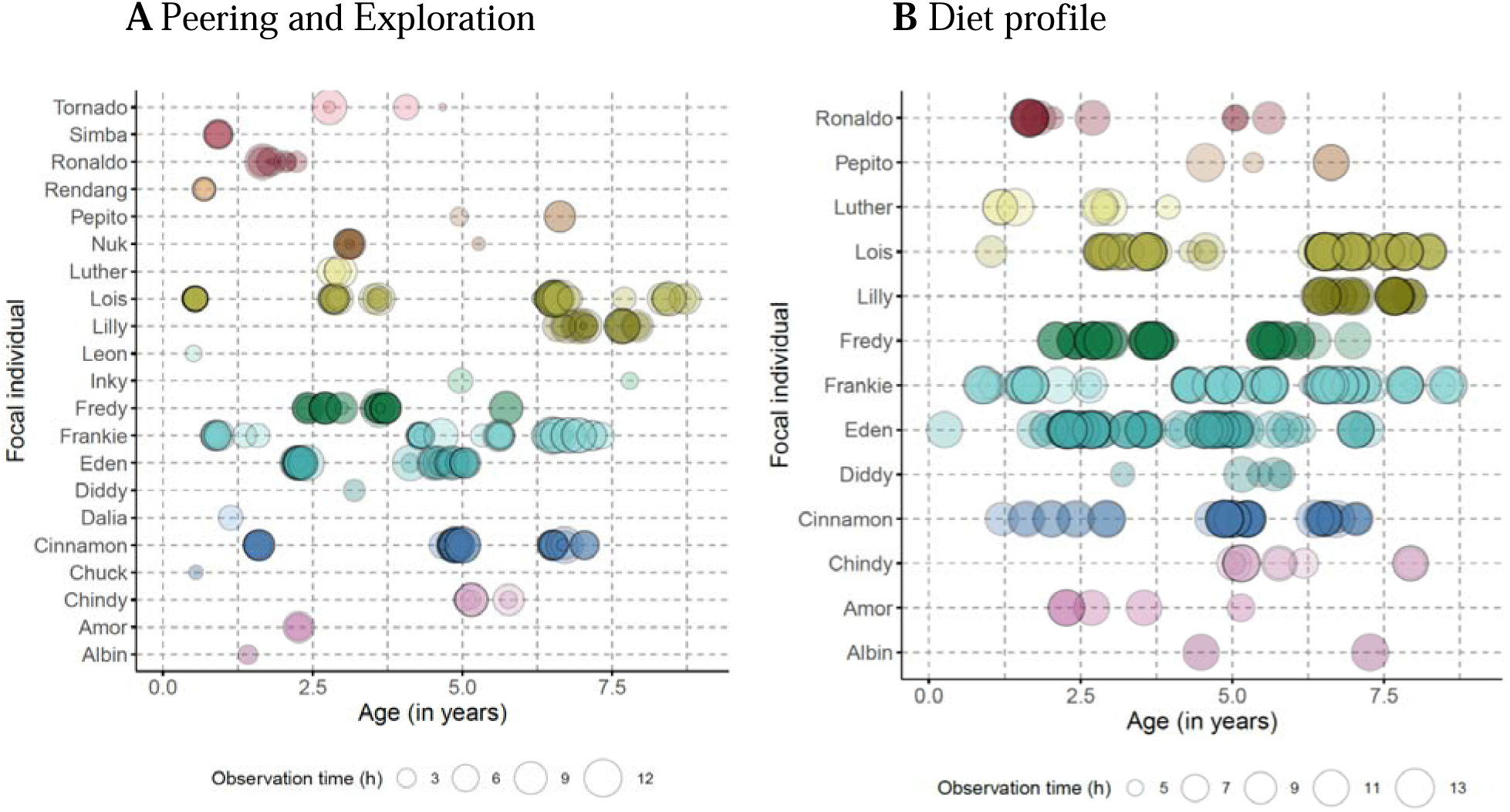
Focal immatures and the ages over which they were sampled. Graphs are for (**A**) peering and exploration behaviors and (**B**) diet profile. Each circle represents a focal follow. The size of the circle corresponds to the duration of the focal follow. The darker the circle, the higher the number of focal follows around that age.

**Fig. S2.**
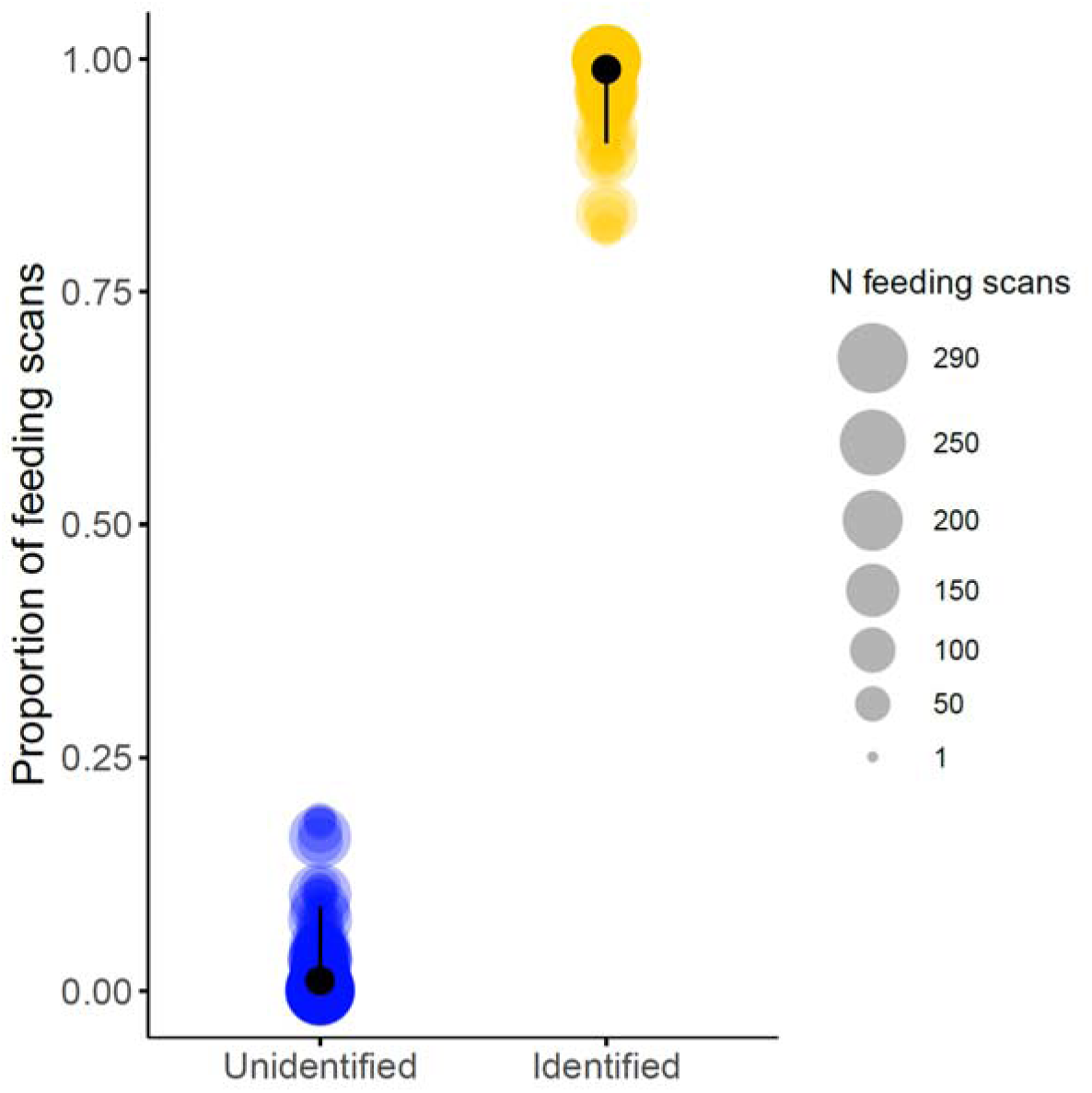
Proportion of feeding scans in which food items were unidentified and identified. Each circle represents a follow, with circle size indicating the total number of feeding scans (feeding scans in which food items were identified + feeding scans in which food items were unidentified) recorded during the follow. Filled, black circles represent the mean proportions of feeding scans in which the food items were unidentified (blue) and identified (yellow). The black lines represent the 95%CI around the means.

**Fig. S3.**
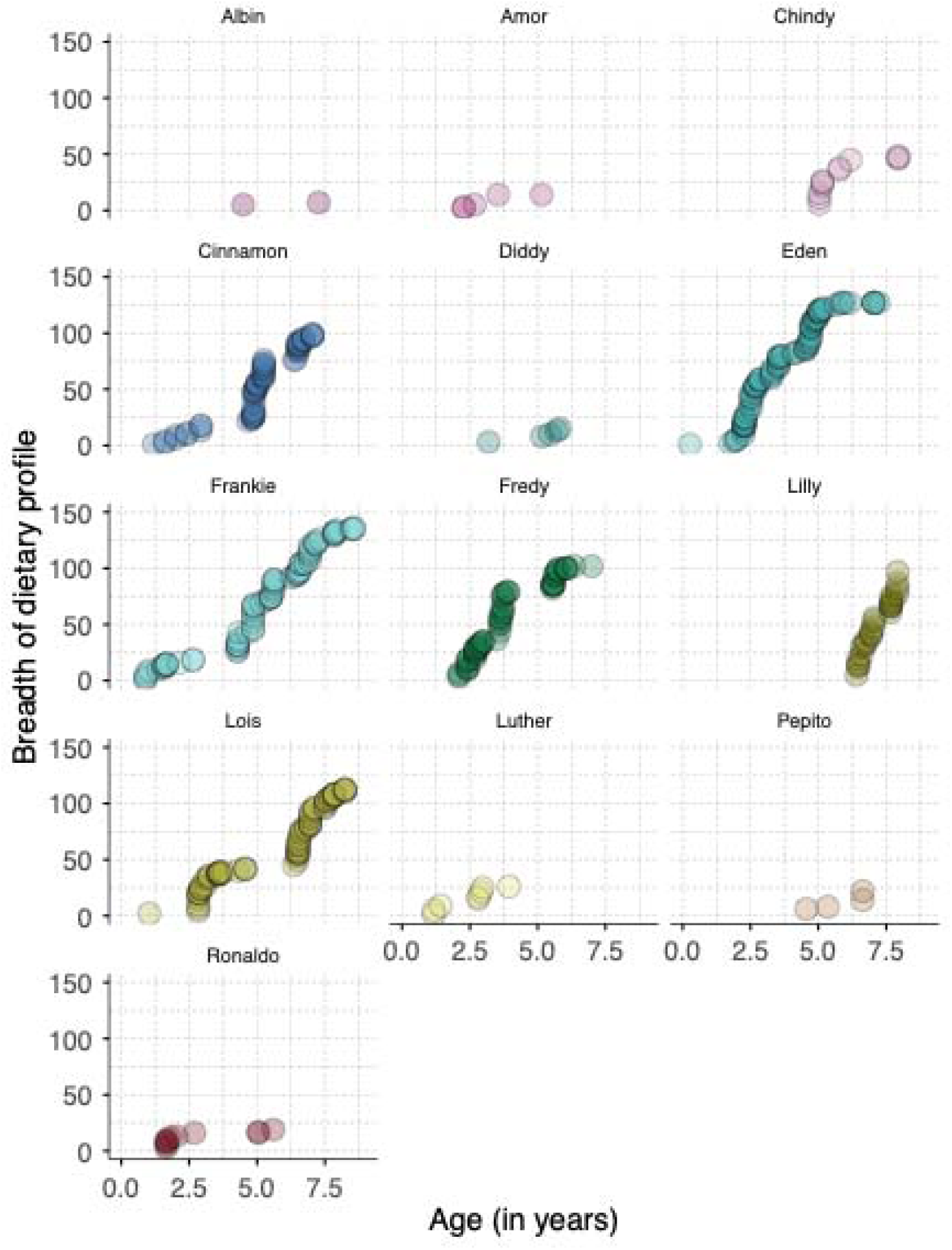
Breadth of the diet profile. Cumulative number of recorded, identified unique food items eaten by 13 focal dependent immatures from birth till the onset of independence at around8.5 years of age. Each circle represents the cumulative count at a specific age for each immature. The darker the circle, the higher the number of focal follows at that age. Note that the breadth of the dietary profile at independence cannot be directly extracted from these graphs. We modelled it via a non-linear (Michaelis Menten) function of age (described in Statistical analysis), using the (plotted) data at the individual level.

**Fig. S4.**
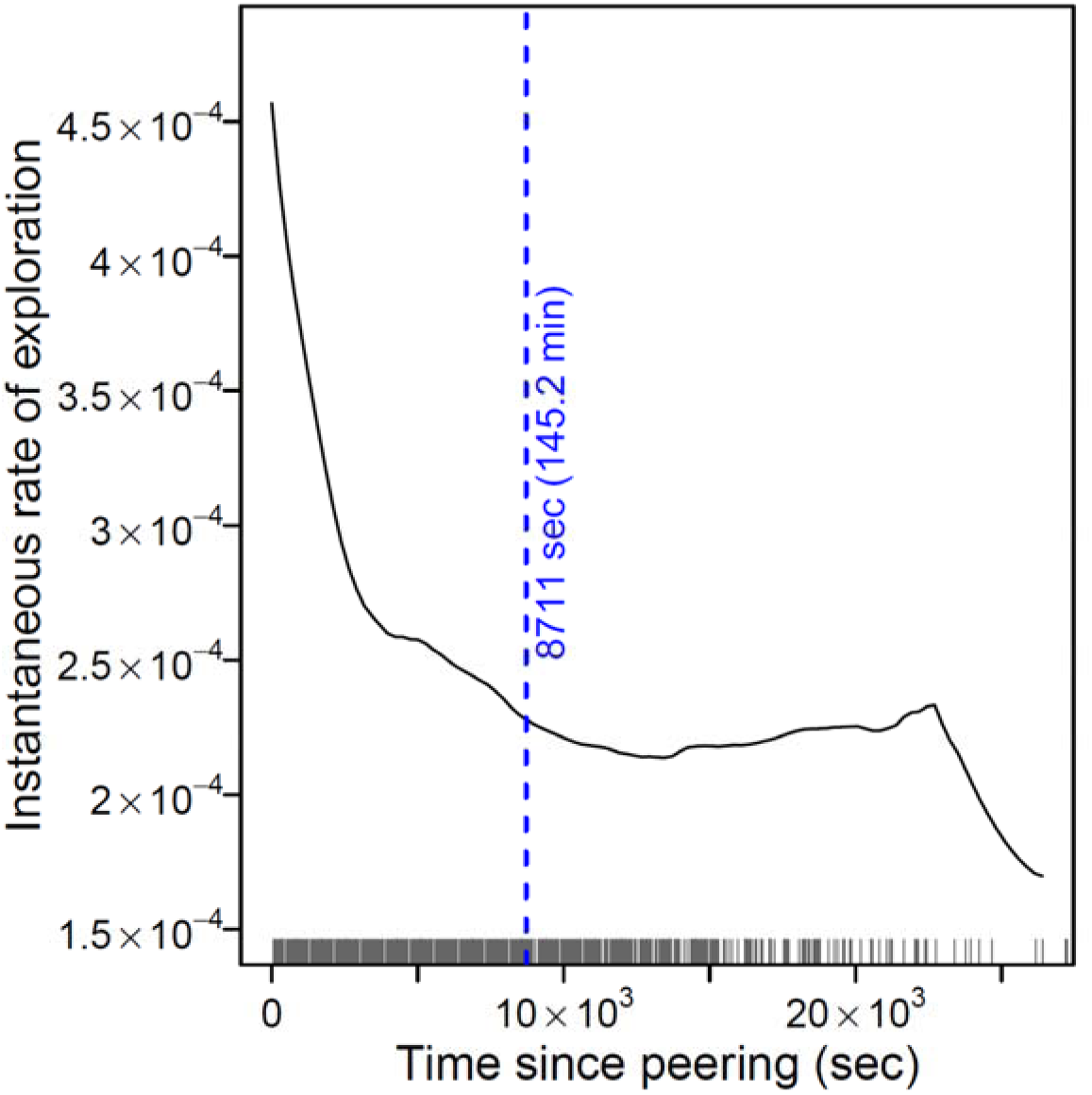
Time to independence of an exploration event following a peering event. Time differences between a peering event and successive exploration events till the next peering event during a focal follow are plotted as grey lines (*N*=4483 time differences). The solid, black curve shows the instantaneous rate of exploration (hazard rate). The dashed blue line marks the timepoint at which the hazard function reaches a near-zero slope. The second dip in hazard rate is a result of very few peering events being separated by around 6 hours.

**Fig. S5.**
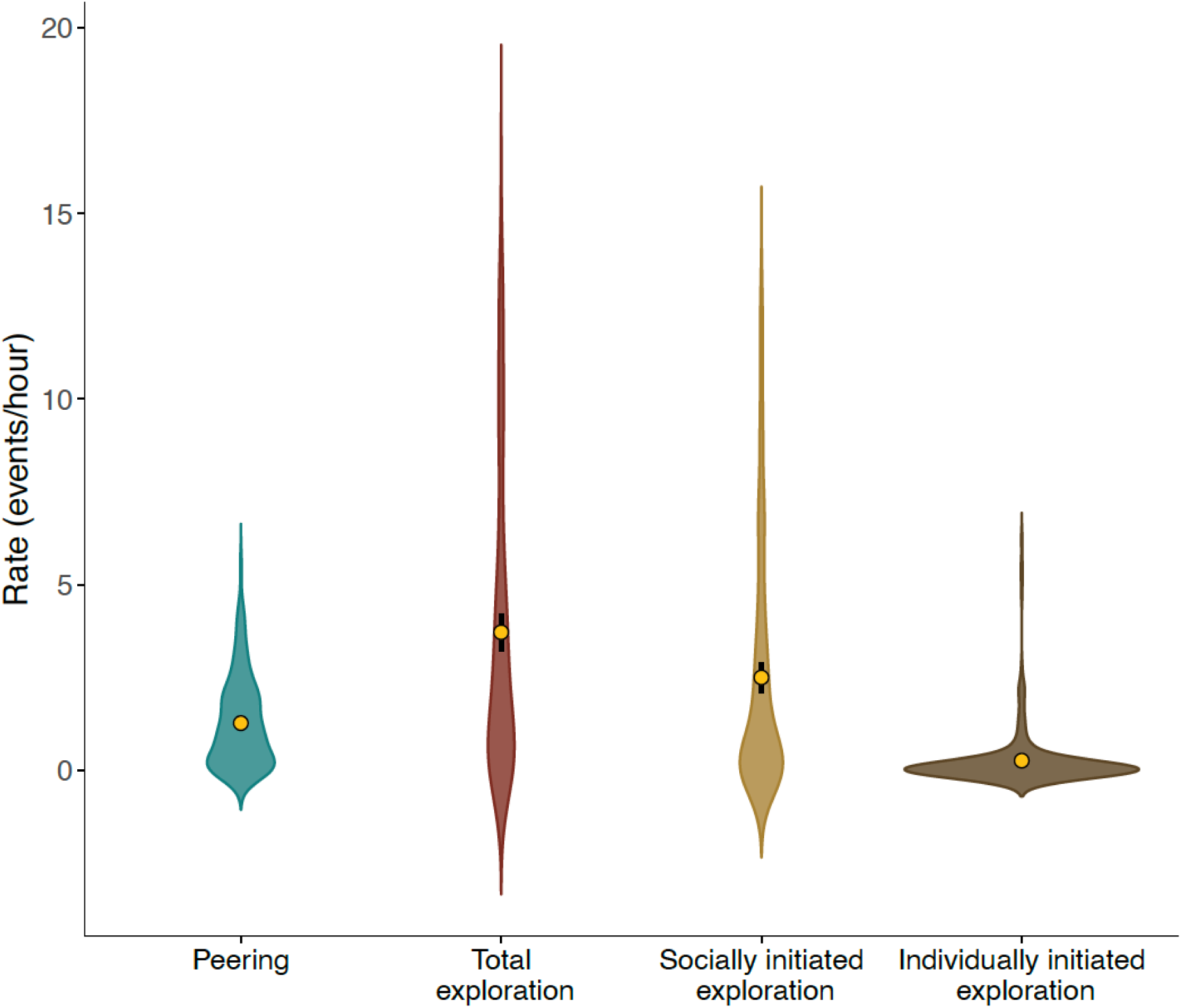
Propensities to seek information socially and individually. Rates of peering, total exploration, socially initiated exploration, and individually initiated exploration are plotted. The filled, yellow circle marks the mean rate over all focal follow (*N*=257) and the thick, black lines represent the 95%CI around the means. Note that the violin plot features a kernel density estimation of the underlying distribution. As a result, the distributions (but not the observed mean or the 95%CI around the mean) include negative rates.

**Fig. S6.**
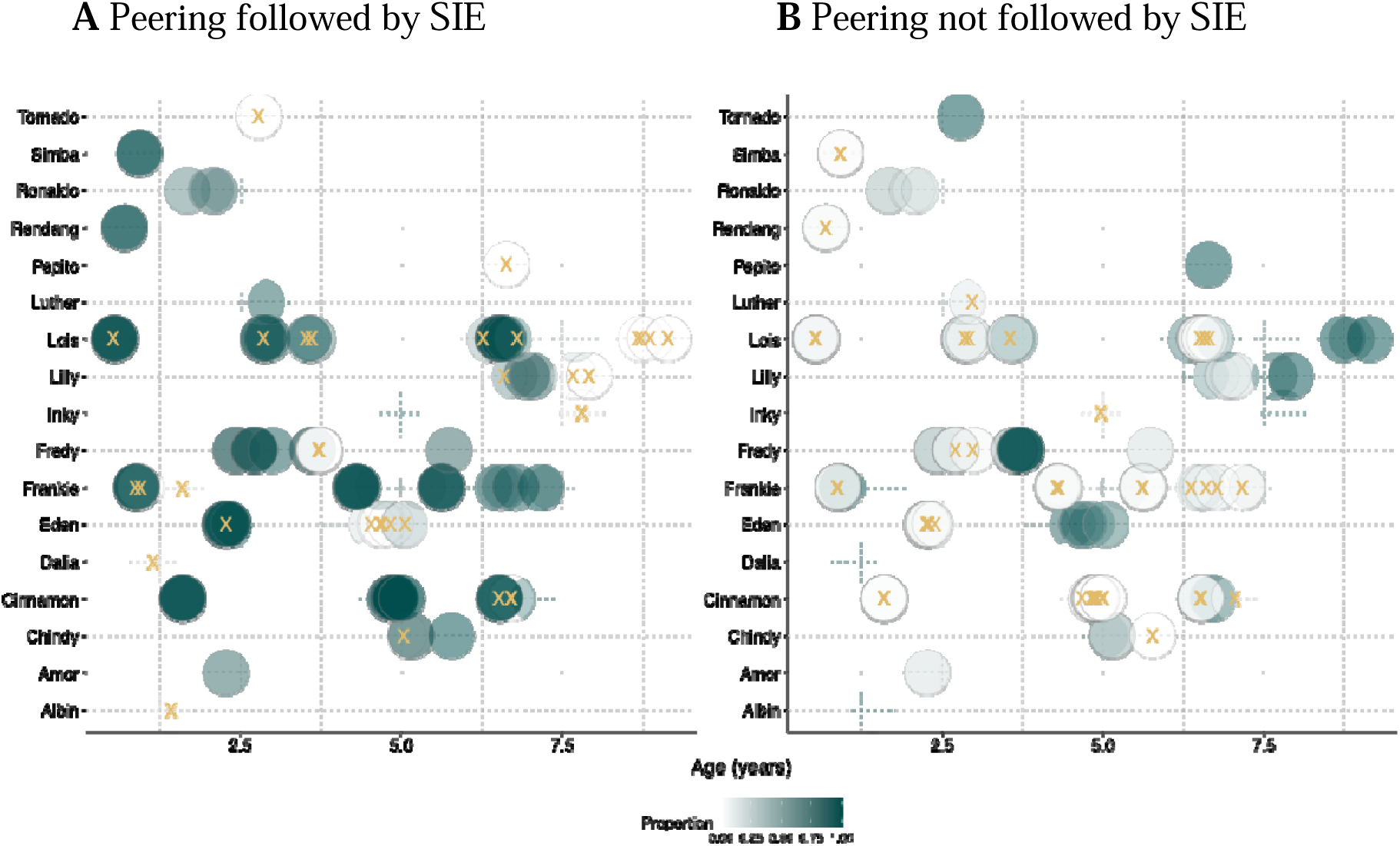
Direct behavioral linkage between peering and socially initiated exploration. The graphs show the proportion of peering events during each focal follow which the focal immature (**A**) followed up with and (**B**) did not follow up with a socially initiated exploration across their dependency period. Each filled circle denotes a focal follow; the shade represents the respective proportion of peering events. To visualize overlapping data points, transparency was added. The yellow ‘X’ indicates a proportion of zero. Note that each follow in (**A**) also appears in (**B**).

**Table S1.**
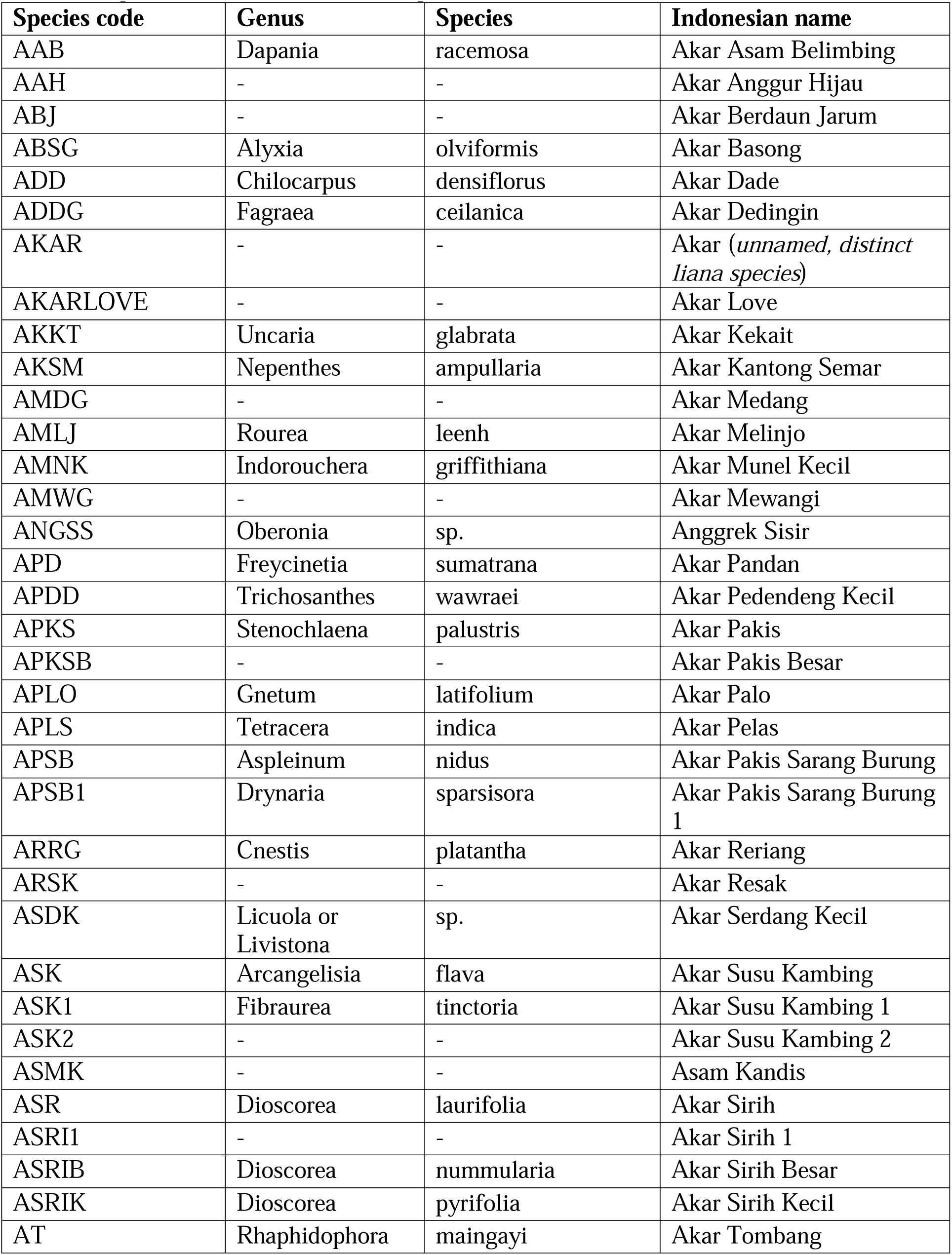
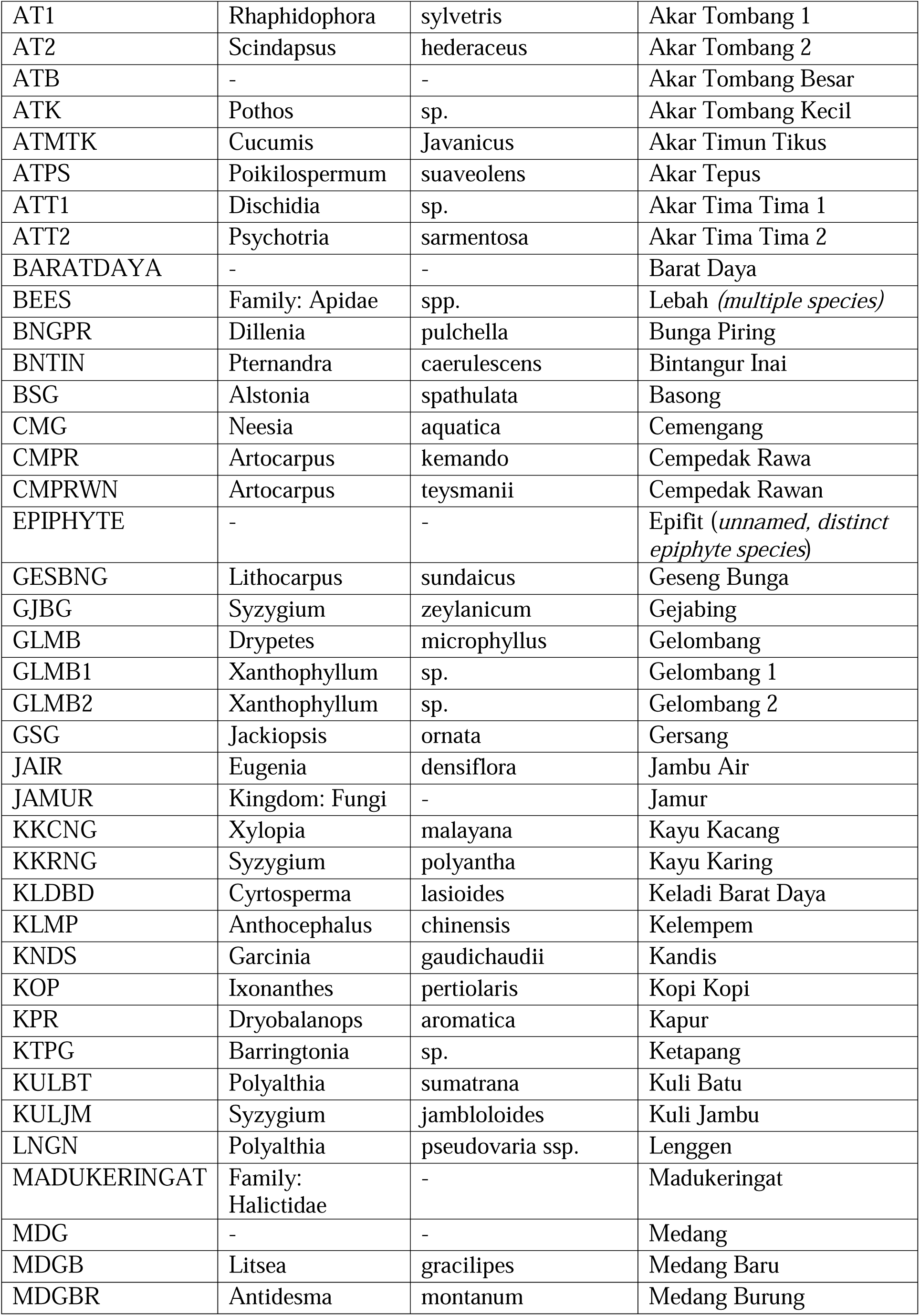
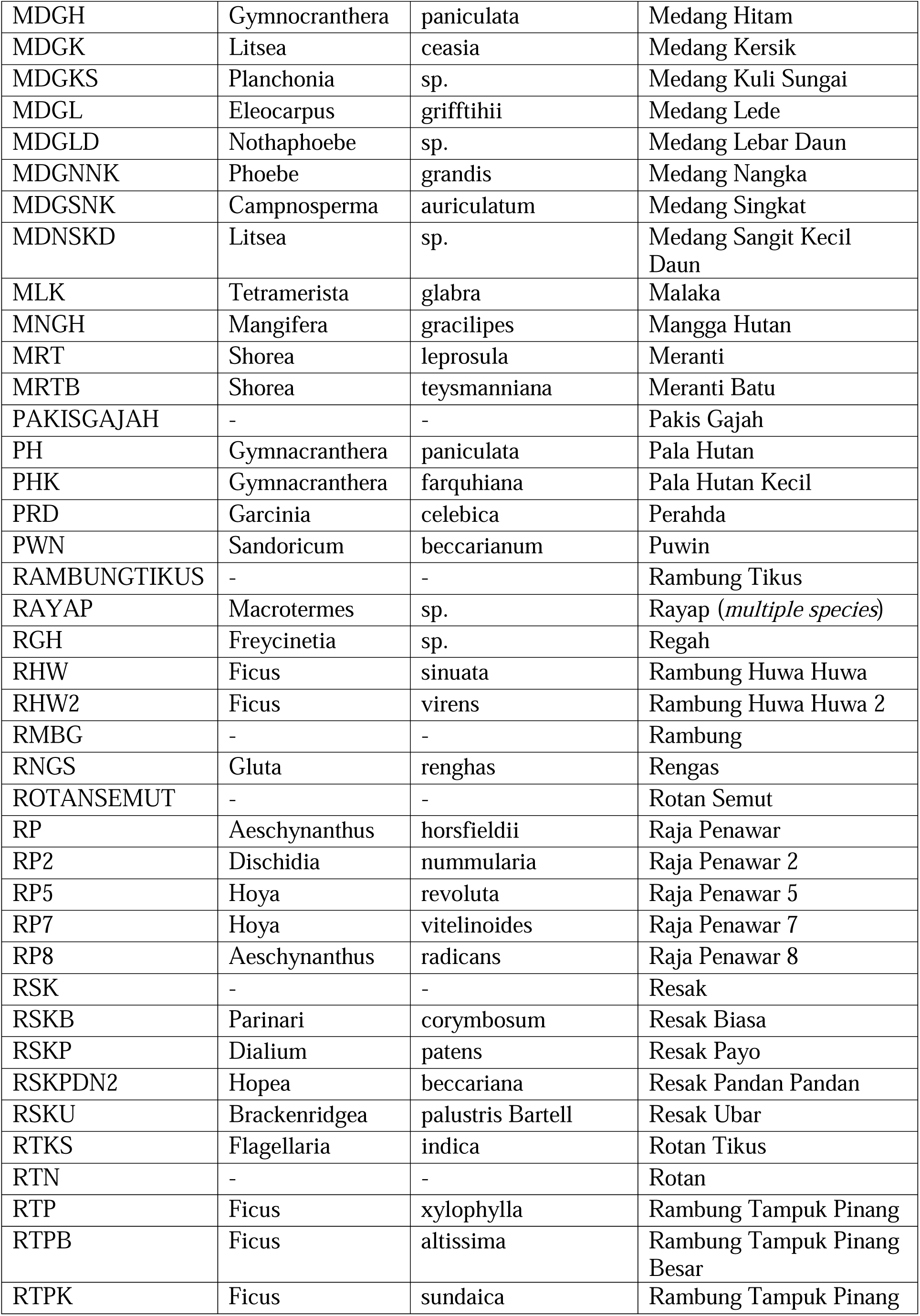
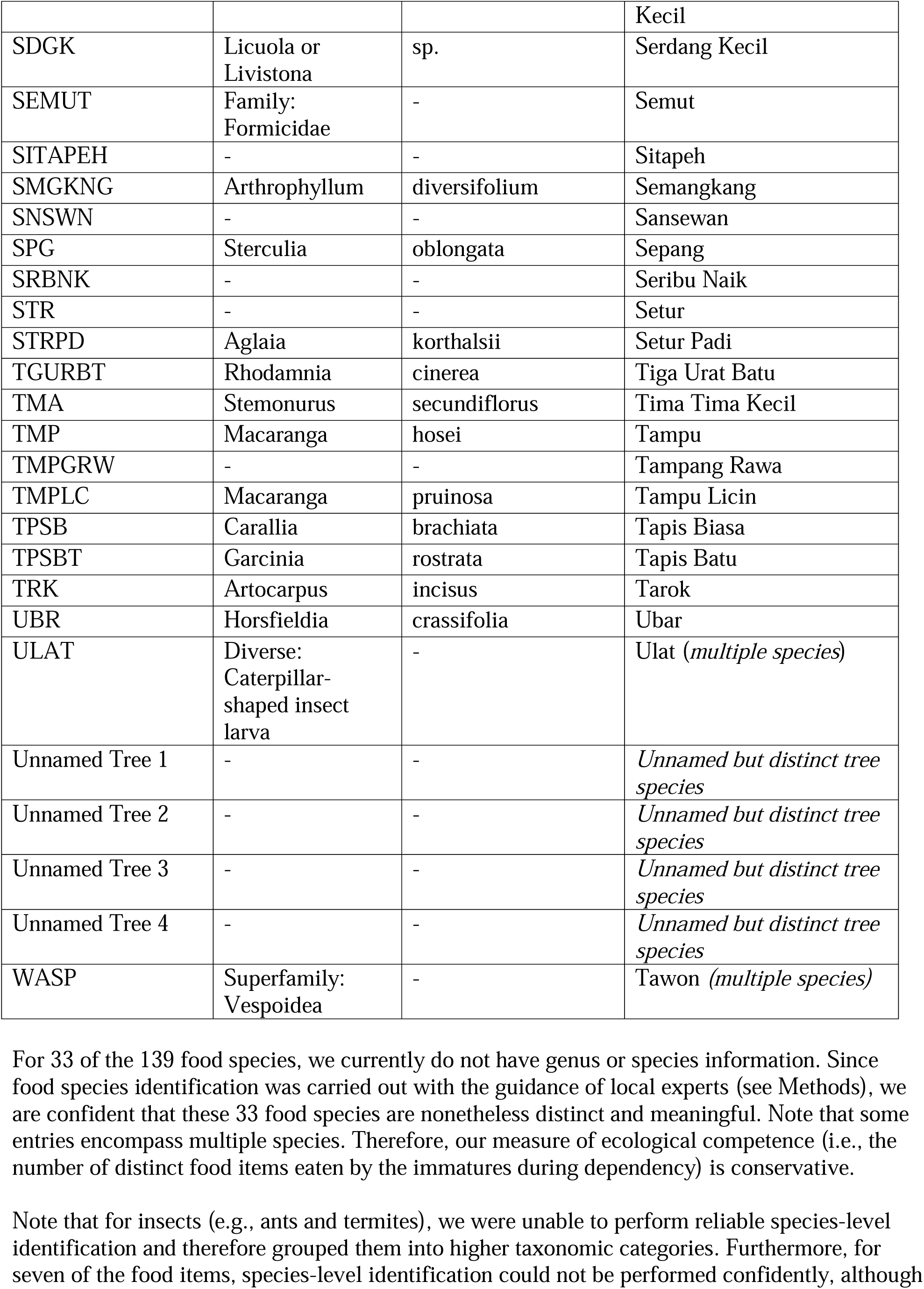
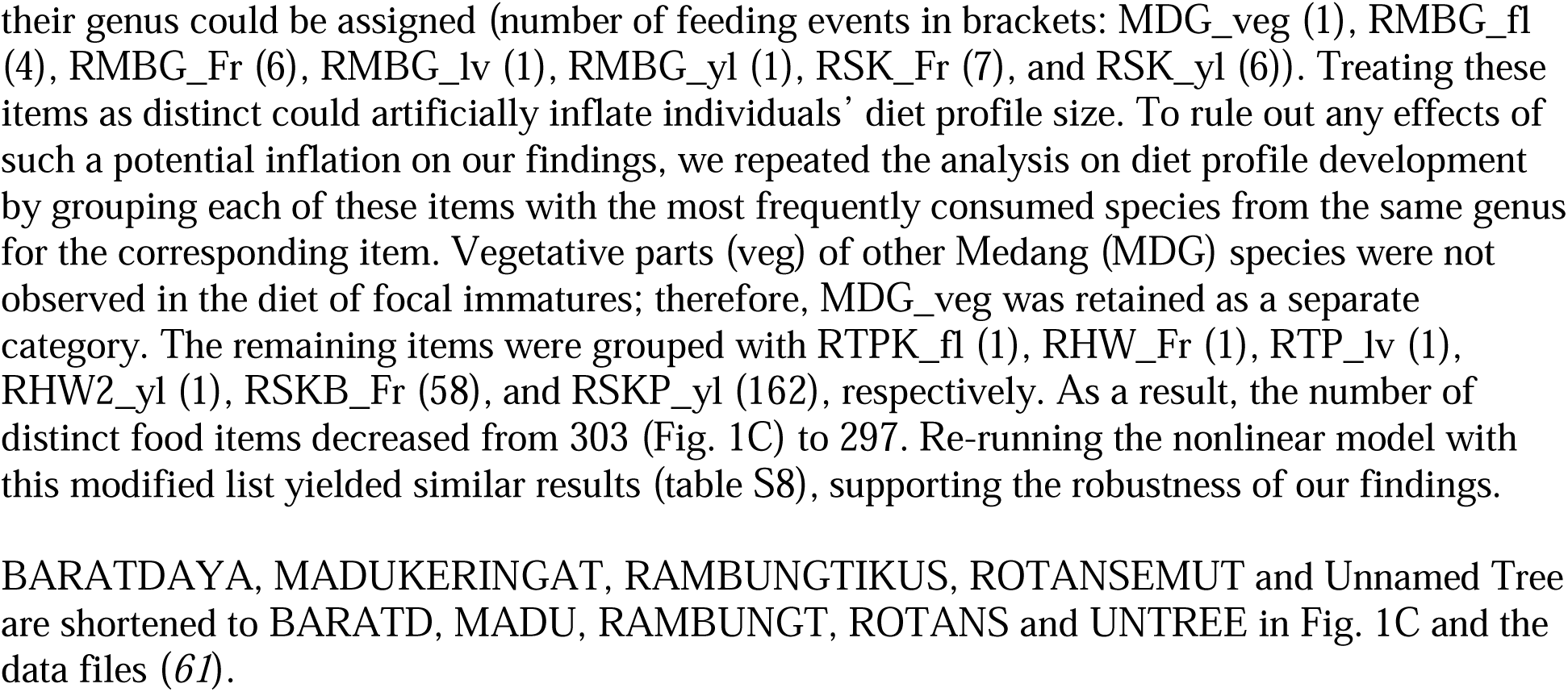
List of food species of focal immature Sumatran orangutans. The species codes of the food items (used in Fig. 1C in the Main text and data files), their genus and species details and the respective Indonesian names are provided here.

**Table S2.** Leave-one-out cross validation. : Do immatures show distinct individual propensities in seeking information socially and individually? Results of the model comparisons based on LOO-CV are shown. Model with a higher predictive performance is marked with an asterisk (*).

| Model | Dataset | Behavior | ΔELPD | ΔSE |
| --- | --- | --- | --- | --- |
| U1* | Total | Peering | 0 | 0 |
| Reduced | Total | Peering | -18.6 | 9.2 |
| U2* | $\leq 145$ | Socially initiated exploration | 0 | 0 |
| Reduced | $\leq 145$ | Socially initiated exploration | -13.5 | 6.0 |
| U3* | $> 145$ | Individually initiated exploration | 0 | 0 |
| Reduced | $> 145$ | Individually initiated exploration | -3.7 | 3.8 |
| U4* | Total | Total exploration | 0 | 0 |
| Reduced | Total | Total exploration | -31.7 | 10.7 |
| SU1* | $> 70$ | Individually initiated exploration | 0 | 0 |
| Reduced | $> 70$ | Individually initiated exploration | -1.5 | 2.5 |
| SU2* | $> 130$ | Individually initiated exploration | 0 | 0 |
| Reduced | $> 130$ | Individually initiated exploration | -1.5 | 3.3 |
| SU3* | $> 160$ | Individually initiated exploration | 0 | 0 |
| Reduced | $> 160$ | Individually initiated exploration | -3.2 | 4.2 |

Leave-one-out cross-validation (LOO-CV) allowed us to examine whether the out-of-sample predictive performance of a model is improved by the addition of model terms (here, the random intercepts and slopes). It does so by estimating the difference in the expected log predictive density (ELPD). $\Delta$ ELPD indicates the ELPD difference between the full and the reduced model, and $\Delta$ SE the corresponding standard error. Note that the full models had higher predictive performance than the reduced models for all the dependent variables. However, for individually initiated exploration, the difference between the full models and the reduced models were small relative to their uncertainty. We nevertheless retained the full model to derive individual propensity to peer and explore because it preserves the full set of covariates and variance structure of interest.

**Table S3.**
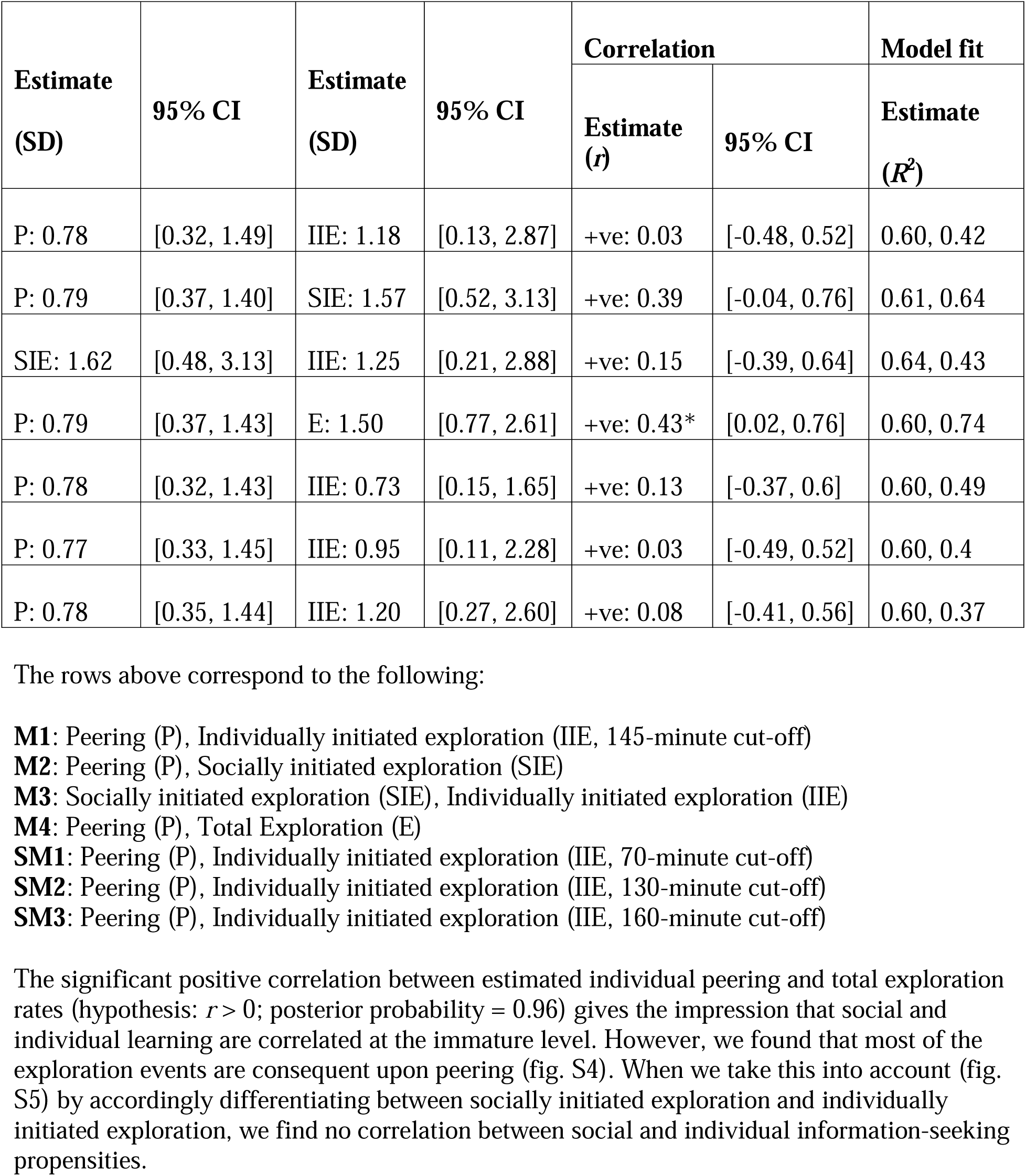
Correlation between information-seeking rates at the individual level using multivariate GLMM (M1-M4, SM1-SM3): Are the social and individual learning propensities (i.e., information-seeking rates) correlated at the immature-level? *indicates significant posterior probability.

**Table S4.** Diet profile development tested using Michaelis-Menten nonlinear, hierarchical Bayesian model (NM1) : Do individuals’ propensities to peer and individually explore (145-minute cut-off) interact to contribute to the development ecological competence? Diet profile was modelled as a Michaelis-Menten function of age. *R*^2^=0.55 [0.49, 0.60], model with a linear interaction term.

| Variable | Estimate | Estimate Error | 95% CI |
| --- | --- | --- | --- |
| <u>Immature identity (Number of levels: 13):</u> |  |  |  |
| SD($V_{\max}$ Intercept) | 52.12 | 5.83 | [41.34, 63.96] |
| SD($K$ Intercept) | 8.20 | 1.29 | [5.72, 10.77] |
| <u>Regression Coefficients:</u> |  |  |  |
| $V_{\max}$ | | | |
| Intercept | 150.01* | 9.06 | [132.12, 168.17] |
| Sex | 14.26* | 4.04 | [6.35, 22.15] |
| Estimated peering | 0.63 | 3.79 | [-6.88, 8.03] |
| Estimated individually initiated exploration | 1.75 | 3.85 | [-5.50, 9.17] |
| Estimated peering*Estimated individually initiated exploration | -36.65* | 16.92 | [-70.11, -3.82] |
| $K$ | | | |
| Intercept | 4.10* | 1.06 | [1.98, 6.17] |
| Sex | 1.05* | 0.50 | [0.06, 2.04] |
| **Base** |  |  |  |
| Intercept | 0.18 | 0.18 | [0.00, 0.65] |
| <u>Further Distributional Parameters:</u> |  |  |  |
| shape | 5.80 | 0.49 | [4.87, 6.81] |

**Table S5.** Diet profile development tested using Michaelis-Menten nonlinear, hierarchical Bayesian model (SNM1, sensitivity analysis) : Do individuals’ propensities to peer and individually explore (70-minute cut-off) interact to contribute to the development ecological competence? Diet profile was modelled as a Michaelis-Menten function of age *R*^2^=0.55 [0.49, 0.60], model with a linear interaction term.

| Variable | Estimate | Estimate Error | 95% CI |
| --- | --- | --- | --- |
| <u>Immature identity (Number of levels: 13):</u> |  |  |  |
| SD($V_{\max}$ Intercept) | 51.44 | 5.91 | [40.62, 63.84] |
| SD($K$ Intercept) | 8.15 | 1.33 | [5.64, 10.92] |
| <u>Regression Coefficients:</u> |  |  |  |
| $V_{\max}$ | | | |
| Intercept | 150.98* | 9.17 | [133.00, 169.40] |
| Sex | 14.19* | 3.88 | [6.73, 21.97] |
| Estimated peering | 0.16 | 3.91 | [-7.50, 7.81] |
| Estimated individually initiated exploration | 1.86 | 3.88 | [-5.92, 9.49] |
| Estimated peering*Estimated individually initiated exploration | -34.41* | 14.10 | [-62.17, -6.29] |
| $K$ | | | |
| Intercept | 4.13* | 1.01 | [2.16, 6.06] |
| Sex | 1.05* | 0.50 | [0.07, 2.05] |
| **Base** |  |  |  |
| Intercept | 0.17 | 0.17 | [0.00, 0.65] |
| <u>Further Distributional Parameters:</u> |  |  |  |
| shape | 5.78 | 0.51 | [4.85, 6.86] |

**Table S6.** Diet profile development tested using Michaelis-Menten nonlinear, hierarchical Bayesian model (SNM2, sensitivity analysis) : Do individuals’ propensities to peer and individually explore (130-minute cut-off) interact to contribute to the development ecological competence? Diet profile was modelled as a Michaelis-Menten function of age. *R*^2^=0.55 [0.49, 0.60], model with a linear interaction term.

| Variable | Estimate | Estimate Error | 95% CI |
| --- | --- | --- | --- |
| <u>Immature identity (Number of levels: 13):</u> |  |  |  |
| SD($V_{\max}$ Intercept) | 52.55 | 5.78 | [41.54, 63.81] |
| SD($K$ Intercept) | 8.32 | 1.29 | [5.82, 10.98] |
| <u>Regression Coefficients:</u> |  |  |  |
| $V_{\max}$ | | | |
| Intercept | 149.38* | 8.86 | [131.61, 167.07] |
| Sex | 14.25* | 4.04 | [6.39, 22.22] |
| Estimated peering | 0.67 | 3.76 | [-6.60, 7.88] |
| Estimated individually initiated exploration | 1.50 | 3.93 | [-6.02, 9.12] |
| Estimated peering*Estimated individually initiated exploration | -39.37* | 16.92 | [-76.69, -2.41] |
| $K$ | | | |
| Intercept | 4.12* | 1.01 | [2.12, 6.08] |
| Sex | 1.04* | 0.52 | [0.08, 2.02] |
| **Base** |  |  |  |
| Intercept | 0.17 | 0.17 | [0.01, 0.61] |
| <u>Further Distributional Parameters:</u> |  |  |  |
| shape | 5.80 | 0.49 | [4.88, 6.81] |

**Table S7.** Diet profile development tested using Michaelis-Menten nonlinear, hierarchical Bayesian model (SNM3, sensitivity analysis) : Do individuals’ propensities to peer and individually explore (160-minute cut-off) interact to contribute to the development ecological competence? Diet profile was modelled as a Michaelis-Menten function of age. *R*^2^=0.55 [0.49, 0.60], model with a linear interaction term.

| **Variable** | **Estimate** | **Estimate Error** | **95% CI** |
| --- | --- | --- | --- |
| <u>Immature identity (Number of levels: 13):</u> |  |  |  |
| SD($V_{\max}$ Intercept) | 50.91 | 5.79 | [39.99, 62.80] |
| SD($K$ Intercept) | 7.90 | 1.30 | [5.53, 10.57] |
| <u>Regression Coefficients:</u> |  |  |  |
| **$V_{\max}$** |  |  |  |
| Intercept | 152.39* | 9.10 | [134.88, 170.25] |
| Sex | 13.97* | 3.96 | [6.17, 21.84] |
| Estimated peering | 0.67 | 3.87 | [-6.90, 8.21] |
| Estimated individually initiated exploration | 2.00 | 3.93 | [-5.74, 9.62] |
| Estimated peering*Estimated individually initiated exploration | -46.04* | 17.24 | [-79.55, -11.78] |
| **$K$** |  |  |  |
| Intercept | 4.13* | 1.02 | [2.15, 6.14] |
| Sex | 1.06* | 0.49 | [0.11, 2.04] |
| **Base** |  |  |  |
| Intercept | 0.18 | 0.17 | [0.01, 0.62] |
| <u>Further Distributional Parameters:</u> |  |  |  |
| shape | 5.76 | 0.51 | [4.82, 6.79] |

**Table S8.** Diet profile development tested using Michaelis-Menten nonlinear, hierarchical Bayesian model (certain food species merged, see table S1) : Do individuals’ propensities to peer and individually explore (145-minute cut-off) interact to contribute to the development ecological competence? Diet profile was modelled as a Michaelis-Menten function of age. *R*^2^=0.55 [0.50, 0.60], model with a linear interaction term.

| Variable | Estimate | Estimate Error | 95% CI |
| --- | --- | --- | --- |
| <u>Immature identity (Number of levels: 13):</u> |  |  |  |
| SD($V_{\max}$ Intercept) | 52.45 | 5.62 | [41.77, 63.91] |
| SD($K$ Intercept) | 8.25 | 1.31 | [5.78, 10.90] |
| <u>Regression Coefficients:</u> |  |  |  |
| $V_{\max}$ | | | |
| Intercept | 149.36* | 9.10 | [131.77, 167.22] |
| Sex | 14.23* | 4.08 | [6.16, 22.35] |
| Estimated peering | 0.61 | 4.00 | [-7.33, 8.48] |
| Estimated individually initiated exploration | 1.88 | 3.93 | [-5.66, 9.54] |
| Estimated peering*Estimated individually initiated exploration | -39.05* | 18.87 | [-75.38, -0.59] |
| $K$ | | | |
| Intercept | 4.12* | 1.01 | [2.12, 6.03] |
| Sex | 1.05* | 0.5 | [0.08, 2.02] |
| **Base** |  |  |  |
| Intercept | 0.17 | 0.18 | [0.00, 0.62] |
| <u>Further Distributional Parameters:</u> |  |  |  |
| shape | 5.79 | 0.52 | [4.85, 6.89] |

